# Senescence triggers intracellular acidification and lysosomal alkalinization via ATP6AP2 attenuation in breast cancer cells

**DOI:** 10.1101/2023.04.08.536098

**Authors:** Wei Li, Kosuke Kawaguchi, Sunao Tanaka, Chenfeng He, Yurina Maeshima, Eiji Suzuki, Masakazu Toi

## Abstract

Several chemotherapy drugs induce the senescence of cancer cells; however, the mechanism underlying intracellular pH dysregulation in senescent cells remains unclear. Adenosine triphosphatase H+ transporting accessory protein 2 (ATP6AP2) plays a critical role in maintaining pH homeostasis in cellular compartments. We here report a new function of ATP6AP2 in senescent breast cancer cells induced by doxorubicin and abemaciclib treatment. ATP6AP2 expression was significantly downregulated in senescent cells, leading to aberrant pH levels that impaired lysosome function and caused immune response changes. The drugs caused cell cycle arrest and proliferation suppression through the upregulation of senescence-related genes. Additionally, senescent cells showed altered inflammatory and immune transcriptional profiles by reprogramming the senescence-associated secretory phenotype. These findings suggest that ATP6AP2-mediated pH regulation during therapy-induced senescence may be linked to immune changes in senescent cancer cells. These findings provide novel insights into understanding the cellular mechanisms underlying the response to anti-cancer drugs.

## Introduction

Breast cancer is highly prevalent and is a significant cause of mortality in women worldwide^1^. Currently, neoadjuvant chemotherapy and adjuvant systemic therapy improve the prognosis and increase the survival of breast cancer patients^2^. Despite advances in breast cancer treatment, many patients develop therapy resistance, leading to an unfavorable prognosis^3^.

Cellular senescence is a physiological stress response in normal and cancer cells, characterized by irreversible cell cycle arrest and permanent proliferation suppression^4,5^. A diversity of factors, including excessive activation of oncogenes, accumulation of stress, metabolic disturbances, irradiation, and chemotherapeutic drugs, can lead to cellular senescence^5–8^. However, the specific role of senescence in cancer development has been controversial. Extensive research has shown that senescence suppresses neoplastic formation and tumor progression by restricting the repair and proliferation mechanisms of impaired cells^9–11^. Similarly, senescence inhibits tumor progression by enhancing the anti-tumor immune response^12,13^. Senescence-induced therapies trigger senescence-associated secretory phenotype (SASP)-dependent vascular remodeling to sensitize the cells to the effects of cytotoxic therapeutic drugs^14^. Recent evidence further established that senescent cells secrete multiple pro-inflammatory factors through the SASP that contribute to immune evasion^15–17^. However, it is unclear whether therapy-induced senescence reprograms the SASP to affect inflammatory and immune responses.

Adenosine triphosphatase H^+^ transporting accessory protein 2 (ATP6AP2), also known as the prorenin receptor, is an accessory subunit protein of vacuolar-type adenosine triphosphatase (V-ATPase)^18–20^. V-ATPases are complex multi-subunit enzymes that function as rotary nanomotors to pump protons and maintain intracellular pH (pH_i_) homeostasis^21–23^. ATP6AP2 is predominantly localized in the lysosome and plasma membrane, and plays a crucial role in energy conservation and acidification of intracellular compartments to support cellular biological activity^24,25^. Moreover, several studies have shown that ATP6AP2 can interact with transforming growth factor (TGF)-β1 and Wnt/β-catenin molecules^26,27^. ATP6AP2 promotes the tumorigenesis and progression of many cancer types^28–30^. However, the function of ATP6AP2 in regulating the pH homeostasis of the intracellular environment in breast cancer is unclear.

Recent studies have suggested impaired pH_i_ as a hallmark of cancer^31–33^. Dysregulated pH_i_ dynamics facilitate various cancer cell behaviors such as cell proliferation, migration, metastasis, evasion of apoptosis, and metabolic adaptation^34^. In addition, an acidified intracellular environment suppresses antibody-dependent cytotoxicity in breast cancer cells^35^. Therefore, linking cellular senescence and intracellular acidification, and further exploring the molecular mechanism may provide new insights for the study of breast cancer treatment. Although senescent cells undergo numerous metabolic reprogramming events resulting in modifications in the intracellular environment, it is unknown whether senescence triggers aberrant changes in pH_i_ and lysosomal pH (pH_L_) in breast cancer and the underlying mechanisms driving these effects.

We here show that intracellular acidification occurs in senescent breast cancer cells, along with an abnormal pH_L_. We thereby hypothesized that a specific molecule drives pH_i_ and pH_L_ aberrations in therapy-induced senescent breast cancer cells. To explore this possibility, we first induced the senescence of breast cancer cells using two cycles of therapy *in vitro* with doxorubicin (Doxo) and abemaciclib (Abe). To further elucidate the effect of therapy-induced senescence in breast cancer cells, we used RNA sequencing (RNA-seq) to reveal transcriptome-wide changes in senescent cells versus control cells. Among the identified differentially expressed genes (DEGs) with downregulated expression after senescence, functional analysis highlighted the role of V-ATPase. Therefore, we further explored the effect of V-ATPase and its specific subunits in the therapy-induced senescence of breast cancer cells. Among these subunits, we demonstrated that ATP6AP2 expression is downregulated in therapy-induced senescent breast cancer cells, which causes intracellular acidification (reduction of pH_i_) and lysosomal alkalinization (elevation of pH_L_). Additionally, the downregulation of ATP6AP2 expression under therapy-induced senescence triggered SASP reprogramming to activate various pro-inflammatory factors that regulate inflammation and the immune response.

## Results

### Doxo and Abe suppress proliferation through cell cycle arrest in breast cancer cells

We exploited therapeutic drugs (Doxo and Abe) to treat breast cancer cells (the human triple-negative breast cancer cell line MDA-MB-231 and the human luminal A subtype breast cancer cell line MCF-7) for 24 h without a robust cytotoxic effect. After the cells grew in a drug-free medium for 48 h, they were re-treated with the therapeutic drugs. Various features of cellular senescence were detected at 24 h, 48 h, 72 h, and 96 h (Fig. 1a). The Cell Cycle Kit-8 (CCK-8) assay showed that Doxo significantly suppressed proliferation in both MDA-MB-231 and MCF-7 cells (Fig. 1b). Abe inhibited proliferation in a time- and dose-dependent manner (Fig. 1c). However, 1000 nM Abe caused massive cell death and 24 h treatment was insufficient to suppress proliferation; thus, this concentration of Abe was deemed to be unsuitable for senescence induction in breast cancer cells.

**Fig. 1.**
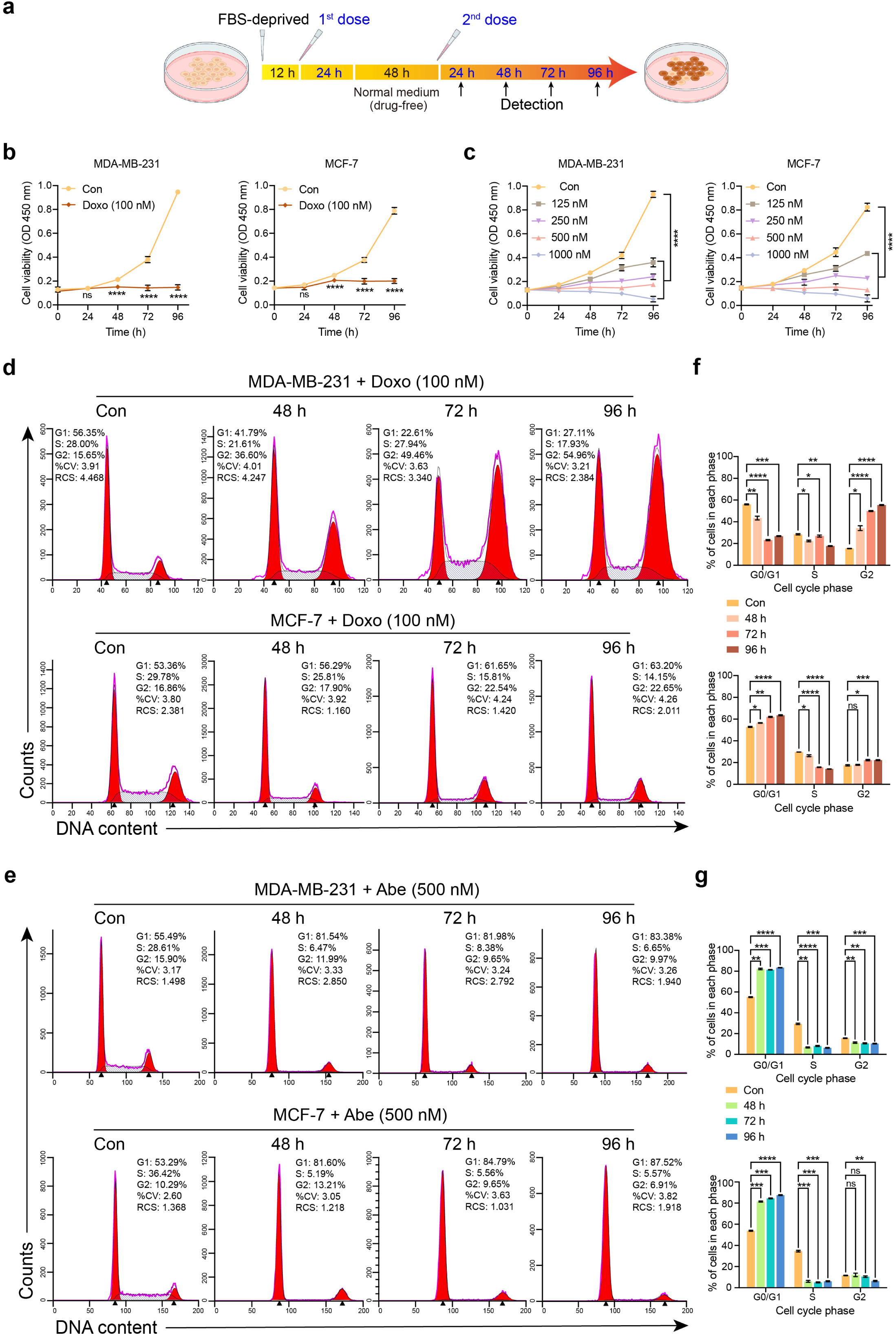
Therapeutic drugs suppress breast cancer cell proliferation through cell cycle arrest. **a** Timeline of cellular senescence induced by therapeutic drugs in breast cancer cells. **b** and **c** Cell proliferation in breast cancer cells treated with doxorubicin (Doxo) at 100 nM (**b**) and abemaciclib (Abe) at different concentrations (up to 1000 nM) (**c**) evaluated by the CCK-8 assay. **d** and **e** Representative images of the cell cycle analysis using propidium iodine (PI) detection by flow cytometry in Doxo (100 nM)-treated (**d**) and Abe (500 nM)-treated (**e**) breast cancer cells. **f** and **g** Quantification of the percentage of cells in each cell cycle phase. Data are shown as means ± SD of three independent experiments. Statistical analyses were performed with two-way ANOVA and Sidak’s multiple-comparisons test **(b**) or Tukey’s multiple-comparisons test (**c**, **f,** and **g**). ns, not significant; \**P* < 0.05, \*\**P* < 0.01, \*\*\**P* < 0.001, \*\*\*\**P* < 0.0001.

Previous studies have demonstrated that DNA damage^36^ and cyclin-dependent kinases (CDKs) inhibition elicit a complex signaling cascade that leads to cell cycle arrest^37,38^. To investigate whether proliferation is suppressed through cell cycle arrest in our treatment conditions, we performed flow cytometry to analyze the cell cycle distribution of breast cancer cells. Doxo elicited G2 cell cycle arrest in MDA-MB-231 cells, in contrast to the G1 phase arrest detected in MCF-7 cells (Fig. 1d and f). Conversely, 500 nM Abe elicited G1 cell cycle arrest in both MDA-MB-231 and MCF-7 cells (Fig. 1e and g), and the cells were also arrested in the G1 phase after 125 nM or 250 nM Abe treatment (Supplementary Fig. 1a–d). These results suggested that therapeutic drugs could substantially inhibit breast cancer cell proliferation, an essential feature of cellular senescence, through cell cycle arrest in a time- and dose-dependent manner.

### Doxo and Abe promote cellular senescence accompanied by elevated p16 and p21 expression in breast cancer cells

We next determined whether Doxo and Abe triggered cellular senescence in breast cancer cells based on senescence-associated β-galactosidase (SA-β-Gal) staining, a widely used senescence marker. Doxo triggered significant cellular senescence after 48 h exposure and showed more pronounced staining with further exposure time, particularly after 96 h of treatment (Fig. 2a and c). Similarly, 500 nM Abe treatment increased the proportion of senescent cells and caused more remarkable cellular senescence at 96 h of exposure (Fig. 2b and d). Although both 250 nM and 125 nM Abe induced cellular senescence, the induction of senescence by 125 nM Abe was not pronounced with less than 50% senescent cells detected (Supplementary Fig. 2a and c), which was lower than the proportion of senescent cells induced in the 250 nM and 500 nM treatment groups (Supplementary Fig. 2b and d). Moreover, the microscope images demonstrated that the cell morphology became irregular and the cell size was enlarged in the senescent cells compared with those of the control cells.

**Fig. 2.**
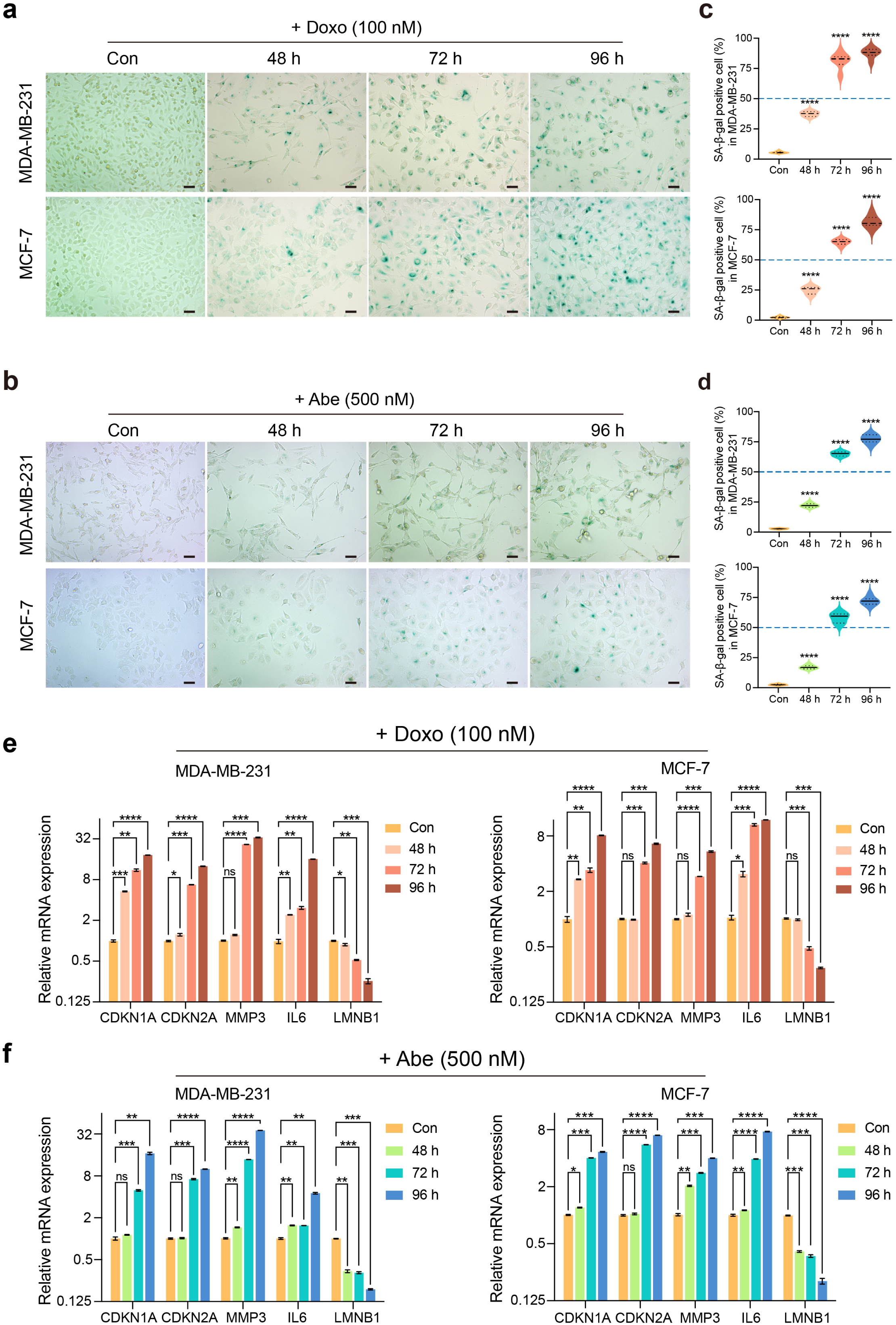
Therapeutic drugs promote cellular senescence with elevated p16 and p21 expression in breast cancer cells. **a** and **b** Representative SA-β-Gal staining images in therapy-challenged MDA-MB-231 and MCF-7 cells treated with doxorubicin (Doxo; 100 nM) (**a**) and abemaciclib (Abe; 500 nM) (**b**). **c** and **d** Quantitative analysis of the percentage of SA-β-Gal–positive cells. At least six separate fields of view were analyzed. **e** and **f** RT-qPCR detection of relative mRNA expression levels of senescence-related genes (*CDKN1A, CDKN2A, MMP3, IL6*, *LMNB1*) in Doxo (100 nM)-treated (**e**) and Abe (500 nM)-treated (**f**) breast cancer cells. Scale bars represent 50 μm. Data are shown as means ± SD of three independent experiments. One-way ANOVA with Dunnett’s multiple-comparisons test (**c** and **d**) and two-way ANOVA with Dunnett’s multiple-comparisons test (**e** and **f**) were performed. ns, not significant; \**P* < 0.05, \*\**P* < 0.01, \*\*\**P* < 0.001, \*\*\*\**P* < 0.0001.

Considering the essential roles of numerous molecular expression profiles correlated with the senescence phenotype, we next sought to verify the molecules considered to be hallmarks of cellular senescence. Reverse transcription-quantitative polymerase chain reaction (RT-qPCR) showed that Doxo simultaneously increased the mRNA levels of *CDKN1A* (p21), *CDKN2A* (p16), matrix metalloproteinase-3 (*MMP3*), and interleukin-6 (*IL6*), accompanied by decreased *LMNB1* levels, with a significant increase in these effects detected at 96 h of treatment (Fig. 2e). In addition, 500 nM Abe elicited significant changes in the senescence-related molecules, with the most prominent effects also observed at 96 h of exposure (Fig. 2f). Moreover, 250 nM Abe treatment yielded molecular changes with a similar trend as found for the 500 nM treatment condition (Supplementary Fig. 2f), whereas the alterations in the expression levels of senescence-associated molecules induced by 125 nM Abe treatment were unstable (Supplementary Fig. 2e). Thus, the 125 nM Abe treatment was excluded from subsequent experiments. Overall, these results demonstrated that therapeutic anti-cancer drugs trigger stable cellular senescence by simultaneously upregulating p21 and p16 expression. Additionally, the senescence phenotype became increasingly more apparent with a prolonged treatment time, particularly under treatment for 96 h.

### Doxo and Abe attenuate ATP6AP2 expression in senescent cells

Given that the therapeutic drugs provoked remarkable cellular senescence after treatment for 96 h, RNA-seq was performed on the therapy-challenged cells for 96 h at 100 nM Doxo and 500 nM Abe, and the transcriptome was compared with that of control cells. The volcano plot in Fig. 3a illustrates the DEGs from the bulk genes expression matrix. Interestingly, in both MDA-MB-231 and MCF-7 cells, the majority of the DEGs were upregulated following Doxo treatment but were downregulated following Abe treatment compared to the expression levels in control cells. Among these DEGs, 29 genes were commonly downregulated in both cell lines, as shown in the Venn diagram in Fig. 3b.

**Fig. 3.**
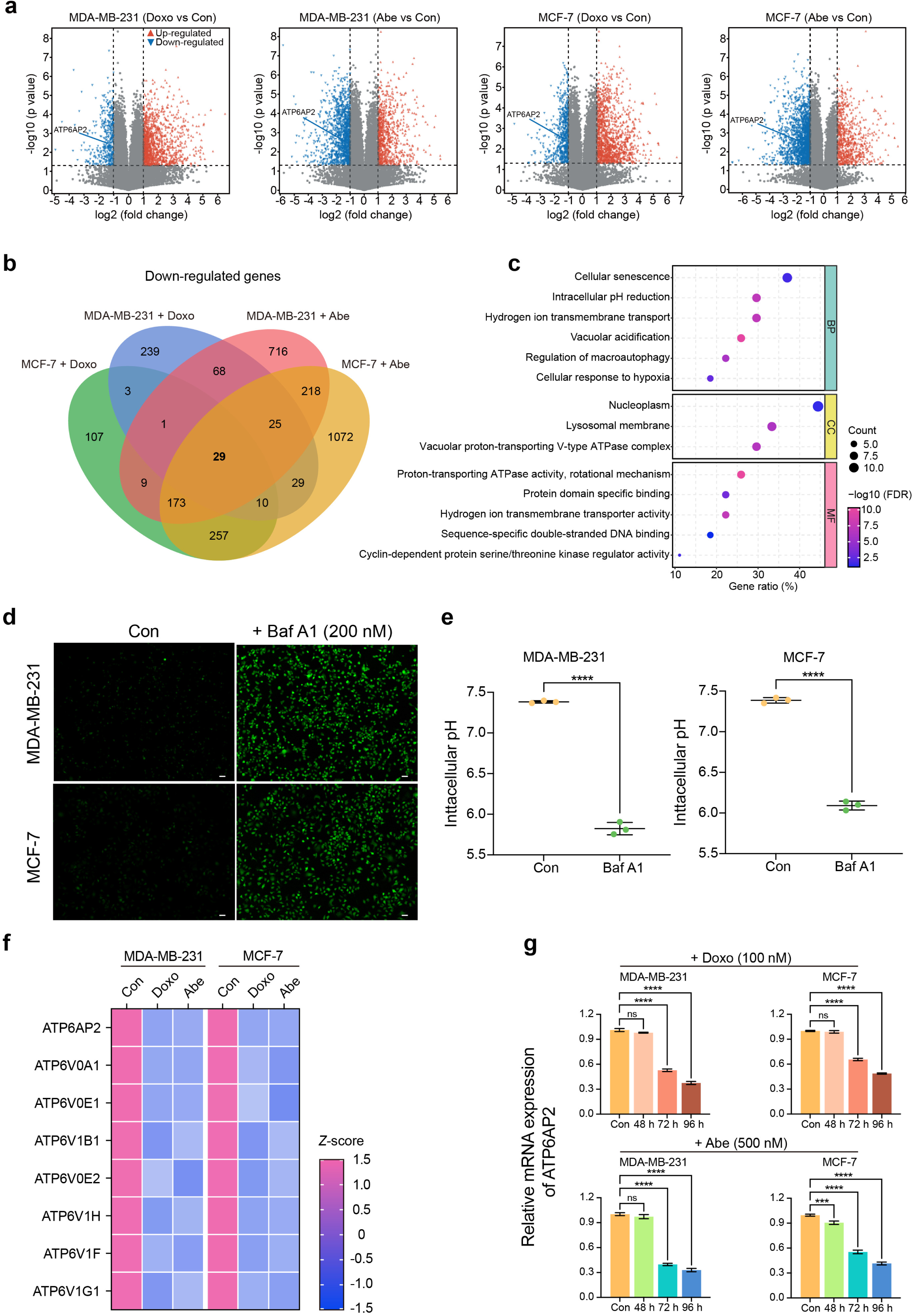
Therapeutic drugs attenuate ATP6AP2 expression in senescent cells. **a** Volcano plot summarizing differentially expressed genes (DEGs) in therapy-challenged cells vs. control cells. Two vertical lines delineate log2 (fold change) of –1 and 1 boundaries, and the horizontal line displays the statistical significance threshold (*p* < 0.05). The blue and red dots indicate down- and upregulated genes, respectively. **b** Four-ellipse Venn diagram outlining overlapping downregulated genes in various therapy-challenged cells. **c** Gene Ontology (GO) enrichment analysis of co-downregulated genes (n = 29) shown with biological process (BP), cellular component (CC), and molecular function (MF) terms. All represented GO terms showed significant enrichment at a false discovery rate (FDR) < 0.05. **d** and **e** Representative fluorescent images of breast cancer cells treated with Bafilomycin A1 (Baf A1, 200 nM) for 3 h stained with pHrodo Green AM (**d**) and quantitative analysis of the pH_i_ (**e**). **f** Heatmap representing the standardized mRNA abundance values (*z*-scores) of the V-ATPase subunits among co-downregulated genes. **g** RT-qPCR detection of relative mRNA expression levels of *ATP6AP2*. Scale bars represent 50 μm. Data are shown as means ± SD of three independent experiments. Data were analyzed by unpaired two-tailed Student’s *t* test (**a** and **e**), Fisher’s exact test with Benjamini-Hochberg multiple-testing correction (**c**), and one-way ANOVA with Dunnett’s multiple-comparisons test (**g**). ns, not significant; \*\*\**P* < 0.001, \*\*\*\**P* < 0.0001.

To explore the biological functions of these downregulated genes, we performed Gene Ontology (GO) and Kyoto Encyclopedia of Genes and Genomes (KEGG) pathway enrichment analyses. GO enrichment analysis revealed that these genes were predominantly enriched in the GO biological processes “cellular senescence” and “intracellular pH reduction,” in the cellular compartment “vacuolar proton-transporting V-type ATPase complex,” and in the molecular function “proton−transporting ATPase activity, rotational mechanism” (Fig. 3c). Pathway enrichment analysis showed that the “cellular senescence,” “cell cycle,” and “lysosome” categories were significantly enriched (Supplementary Fig. 3a). We therefore hypothesized that therapy-induced cellular senescence caused a decrease in V-ATPase, which consequently contributed to pH_i_ decline and abnormal lysosomal function.

To ascertain whether V-ATPase attenuated the therapy-triggered pH_i_ decrease in breast cancer cells, we used 200 nM bafilomycin A1 (Baf A1), a specific V-ATPase inhibitor, to treat MDA-MB-231 and MCF-7 cells over a time gradient. After 1 h of 200 nM Baf A1 treatment, the pH_i_ was lower than that of control cells, and a significant difference was found after 3 h of inhibitor treatment (Supplementary Fig. 3b). Fluorescence microscopy confirmed the pH_i_ decrease under the same treatment condition (Fig. 3d and e). However, 200 nM Baf A1 treatment had no effect on cell viability (Supplementary Fig. 3c).

Subsequently, to elucidate which subunit of V-ATPase was downregulated in senescent cells by treatment with the therapeutic drugs, we first normalized the RNA-seq results of representative subunits of V-ATPase among the 29 downregulated genes, as represented by the heatmap in Fig. 3f and Supplementary Table 1. We then used RT-qPCR to verify the expression levels of the top four downregulated V-ATPase subunits (*ATP6AP2, ATP6V0A1, ATP6V0E1*, and *ATP6V1B1*) compared with those of control (untreated) cells (Supplementary Fig. 3d and e). Although all genes showed a trend of declining expression levels in the therapy-treated cells, only *ATP6AP2* expression remained stably downregulated in Doxo-induced senescent cells at 72 h and 96 h (Fig. 3g, upper). Treatment of Abe at either a 500 nM or 250 nM concentration significantly suppressed *ATP6AP2* expression at 72 h and 96 h in MDA-MB-231 cells, whereas the *ATP6AP2* expression level was significantly reduced in MCF-7 cells at 48 h, 72 h, and 96 h under 250 nM and 500 nM Abe exposure (Fig. 3g, lower; Supplementary Fig. 3f). These results suggested that therapy-induced senescence caused ATP6AP2 suppression.

### Doxo and Abe trigger substantial intracellular acidification in senescent cells

To further prove whether therapeutic drugs induce intracellular acidification in senescent breast cancer cells, we performed pH_i_ detection using pHrodo Green AM staining, a pH-sensitive fluorescent probe, at different time points. Doxo elicited intracellular acidification after 72 h exposure, and the effect was most pronounced at 96 h (Fig. 4a–d). Similarly, 500 nM Abe generated significant intracellular acidification in MDA-MB-231 cells at 72 h and particularly at 96 h (Fig. 4e upper and Fig. 4f). Moreover, treatment of MCF-7 cells with 500 nM Abe not only resulted in marked intracellular acidification at 72 h and 96 h exposure but also induced a certain extent of pH_i_ decrease at 48 h exposure (Fig. 4e lower and Fig. 4g). Likewise, treatment with 250 nM Abe triggered intracellular acidification at 72 h and 96 h exposure (Supplementary Fig. 4a–d).

**Fig. 4.**
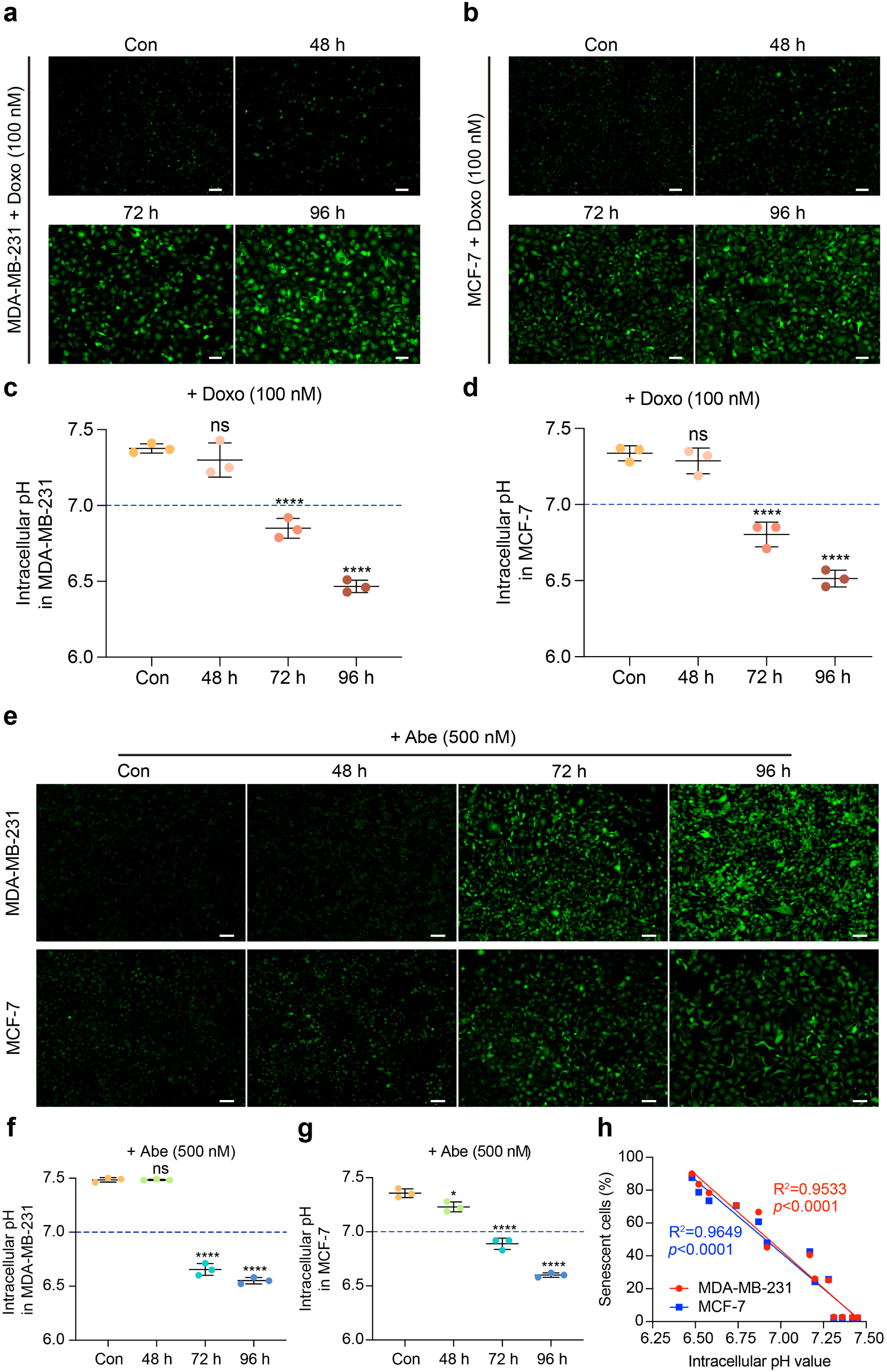
Therapeutic drugs trigger substantial intracellular acidification in senescent cells. a and b Representative fluorescent images of doxorubicin (Doxo; 100 nM)-treated MDA-MB-231 (**a**) and MCF-7 (b) cells stained with pHrodo Green AM. **c** and **d** Quantitative analysis of the intracellular pH in each cell line. **e** Representative fluorescent images of abemaciclib (Abe; 500 nM)-treated MDA-MB-231 (upper panel) and MCF-7 (lower panel) cells stained with pHrodo Green AM. **f** and **g** Quantitative analysis of the intracellular pH in each cell line. **h** Scatterplot illustrating the correlation between the percentage of senescent cells and pH_i_ value in breast cancer cells. Scale bars represent 50 μm. Data are shown as means ± SD of three independent experiments. One-way ANOVA with Dunnett’s multiple-comparisons test (**c, d**, **f**, and **g**) and simple linear regression with Pearson correlation analysis (**h**) were performed. ns, not significant; \**P* < 0.05, \*\*\*\**P* < 0.0001.

Furthermore, we assessed the proportion of senescent cells and the corresponding pH_i_ values under the same treatment conditions to better understand the correlation. Strong positive correlations were found between senescence and pH_i_ in breast cancer cells (Fig. 4h). Moreover, cells with size and morphology changes showed relatively higher fluorescence intensity. These results implied that therapeutic drugs could induce substantial intracellular acidification and a more marked decrease in pH_i_ corresponding with a greater effect on senescence induction.

### Senescence-induced ATP6AP2 attenuation elicits an alkalinized pH_L_ to cause lysosomal impairment

Since the RNA-seq data identified that DEGs in therapy-induced senescent cells were enriched in the signaling pathways of the lysosome and phagosome (Supplementary Fig. 3a), to further clarify whether the therapy-induced impairment of ATP6AP2 can affect pH_L_ in senescent cells, we detected the pH_L_ status by the LysoSensor Yellow/Blue DND-160 probe. For these experiments, the cells were subjected to 72 h and 96 h drug exposure, given that the most significant inhibition of ATP6AP2 and remarkable intracellular acidification were observed under these conditions. Doxo contributed to a significant decrease in yellow fluorescence [emission (Em) 535 nm] intensity with a corresponding increase in blue fluorescence (Em 440 nm) intensity, which caused a substantial reduction in the ratio of yellow to blue fluorescence intensity in MDA-MB-231 cells (Fig. 5a and b). These results suggested that Doxo increased the pH_L_ in MDA-MB-231 cells with 72 h and 96 h treatment. Similarly, Doxo evoked pH_L_ elevation in MCF-7 cells with both 72 h and 96 h exposure (Fig. 5c and d). The pH_L_ similarly increased (based on a decrease in the yellow/blue ratio) in breast cancer cells treated with 500 nM Abe. However, in contrast to the effects of Doxo treatment, the 500 nM Abe treatment appeared to enhance the blue fluorescence rather than weaken the yellow fluorescence as typically found in MDA-MB-231 and MCF-7 cells (Fig. 5e–h). Similarly, pH_L_ increased significantly in MCF-7 cells treated with 250 nM Abe but was only significantly increased after 96 h treatment in MDA-MB-231 cells (Supplementary Fig. 5a and b).

**Fig. 5.**
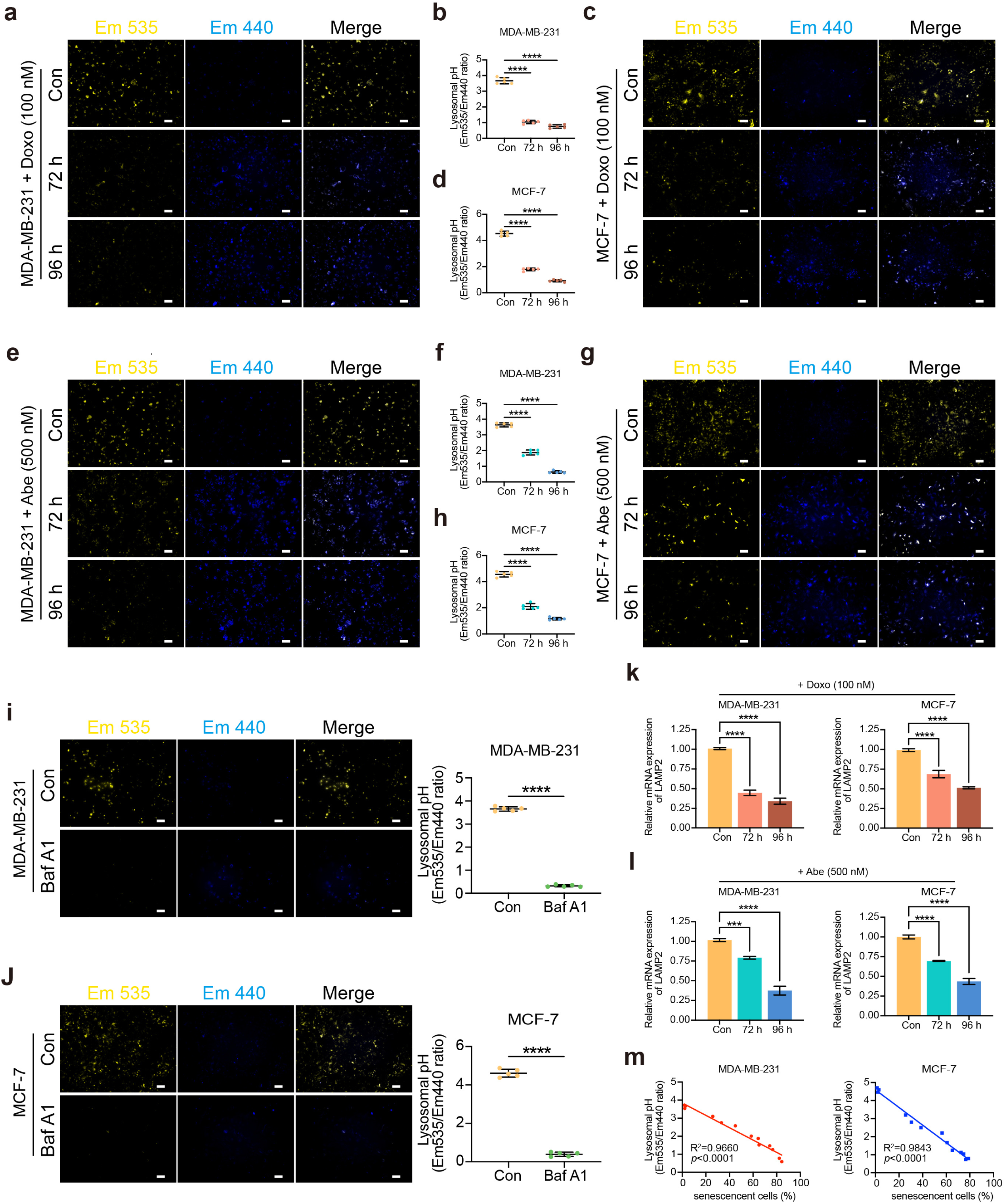
ATP6AP2 attenuation elicits lysosomal pH alkalinization causing lysosomal impairment. a–d Representative fluorescent images of doxorubicin (Doxo; 100 nM)-treated MDA-MB-231 (**a**) and MCF-7 (c) cells stained with LysoSensor Yellow/Blue-DND-160 to assess pH_L_ status. Emission (Em) 535 nm: yellow; Em 440 nm: blue. The pH_L_ status was determined ratiometrically in MDA-MB-231 (n = 5) (**b**) and MCF-7 (n = 5) (**d**) cells. **e–h** Representative fluorescent images of abemaciclib (Abe; 500 nM)-treated MDA-MB-231 (**e**) and MCF-7 (**g**) cells stained with LysoSensor Yellow/Blue-DND-160 to assess pH_L_ status. The pH_L_ status was determined ratiometrically in MDA-MB-231 (n = 5) (**f**) and MCF-7 (n = 5) (**h**) cells. **i** and **j** Representative fluorescent images and quantitative analysis of pH_L_ in Bafilomycin A1 (Baf A1, 200 nM)-treated MDA-MB-231 (**i**) and MCF-7 (**j**) cells stained with LysoSensor Yellow/Blue-DND-160. **k** and **l** RT-qPCR to determine the relative mRNA expression levels of *LAMP2* in each cell line. **m** Scatterplot illustrating the correlation between pH_L_ and the percentage of senescent cells. Scale bars represent 50 μm. Data are shown as means ± SD of three independent experiments. One-way ANOVA with Dunnett’s multiple-comparisons test (**b** and **d**, **f** and **h**, **k** and **l**), unpaired two-tailed Student’s *t* test (**i** and **j**, left panels), and simple linear regression with Pearson correlation (**m**) were performed. \*\*\**P* < 0.001, \*\*\*\**P* < 0.0001.

To further elucidate whether ATP6AP2 attenuated the drug-induced increase in pH_L_, we assayed pH_L_ in breast cancer cells treated with 200 nM Baf A1 for 3 h. As shown in Fig. 5i, the pH_L_ was dramatically increased in MDA-MB-231 cells with Baf A1 exposure and a similar result was observed in Baf A1-treated MCF-7 cells (Fig. 5j). Specifically, compared with the effects of the Doxo and Abe treatments, Baf A1 treatment resulted in strongly reduced yellow fluorescence, along with slightly increased blue fluorescence and a considerably diminished number of lysosomes in the field of view.

Considering that lysosomes are acidic membrane-bound organelles that depend on the acidic pH environment in the lumen of the lysosome to maintain normal function for biological processes^39,40^, we next determined whether the therapy-induced aberrant pH_L_ could generate lysosomal dysfunction. The mRNA expression level of lysosomal-associated membrane protein 2 (*LAMP2*), the main membrane protein of lysosomes^41,42^, was significantly suppressed in breast cancer cells after Doxo treatment at both 72 h and 96 h (Fig. 5k). Similarly, 500 nM Abe also downregulated *LAMP2* expression (Fig. 5l). In contrast, 250 nM Abe only significantly decreased *LAMP2* expression at 96 h of treatment in MDA-MB-231 cells, but at both 72 h and 96 h exposure times in MCF-7 cells (Supplementary Fig. 5c). In addition, RNA-seq data confirmed that the therapeutic drugs downregulated lysosome-related genes compared with those of control cells (Supplementary Fig. 5d and Supplementary Table 2). Notably, these results revealed that the decrease in *LAMP2* expression strongly correlated with the pH_L_ status under the same treatment condition.

To better understand the correlation between therapy-induced cellular senescence and pH_L_ status, we determined the proportion of senescent cells and the corresponding pH_L_ in MDA-MB-231 and MCF-7 cells under the same treatment conditions, respectively. A strong correlation was found between cellular senescence and pH_L_ impairment, suggesting a functional link between the two phenotypes (Fig. 5m). Thus, we concluded that therapy-induced senescence caused ATP6AP2 attenuation, which resulted in an alkalinized pH_L_, and this abnormal pH_L_ environment together with LAMP2 suppression triggered lysosomal impairment.

### Doxo and Abe induce SASP reprogramming to alter inflammatory and immune profiles in breast cancer cells

Cellular senescence has been shown to modify the immune status of the tumor microenvironment through the secretion of multiple inflammatory cytokines/chemokines and immune modulators^43,44^. Therefore, we further explored the impact of therapy-induced senescence on immune and inflammatory processes in breast cancer cells based on the upregulated DEGs identified in RNA-seq analysis. The Venn diagrams in Fig. 6a highlight the significant intersection of 66 upregulated genes among the four groups of MDA-MB-231 and MCF-7 cells treated with Doxo and Abe, respectively.

**Fig. 6.**
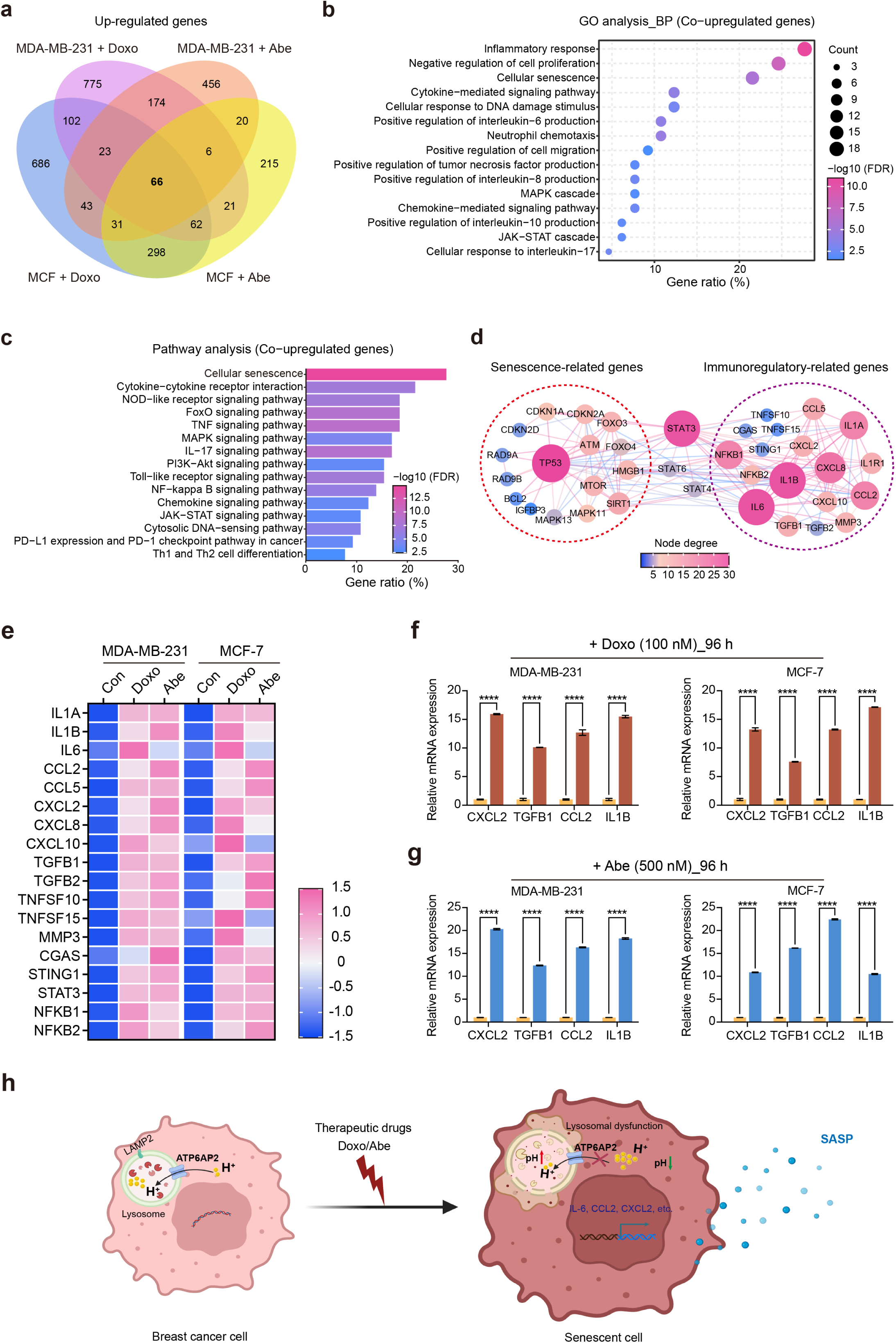
Therapy-induced senescence reprogrammed the senescence-associated secretory phenotype (SASP) to alter inflammatory and immune profiles in breast cancer cells. a Four-ellipse Venn diagram outlining overlapping upregulated genes in various therapy-treated cells. b Gene Ontology (GO) enrichment analysis of co-upregulated genes (n = 66). All represented GO terms with false discovery rate (FDR) < 0.05 are shown. c Pathway enrichment analysis of co-upregulated genes (n = 66). All represented pathway terms with FDR < 0.05 are shown. d STRING network representation of the protein–protein interactions in senescence-related genes (red circles) and immunoregulatory-related genes (purple circles) belonging to co-upregulated genes (FDR < 0.05, interaction score ≥ 0.7). The degree of nodes is represented by size and color. e Heatmap representing the standardized mRNA abundance values (*z*-scores) of the immunoregulatory-related genes. f and g RT-qPCR to detect the relative mRNA expression levels associated with the SASP (*CXCL2, TGFB1, CCL2, IL1B*), respectively. Data are shown as means ± SD of three independent experiments. Two-way ANOVA with Sidak’s multiple-comparisons test (f and g) was performed. \*\*\*\**P* < 0.0001. h Schematic diagram representing the mechanism of ATP6AP2 in therapy-induced senescence and immune regulation via SASP reprogramming in breast cancer cells.

GO enrichment analysis showed that these 66 genes belonged to multiple biological process categories associated with the inflammatory response, cellular senescence, cytokine/chemokine-mediated pathways, JAK-STAT cascade, and positive regulation of IL-6/IL-8/IL-10/tumor necrosis factor (TNF) production, which are known pivotal mechanisms of immune regulation (Fig. 6b). KEGG pathway analysis further suggested that the co-upregulated genes were enriched in several critical immune-related signaling pathways, including cytokine–cytokine receptor interactions, TNF signaling, NF-κB signaling, chemokine signaling, and programmed death-ligand 1 (PD-L1) checkpoint pathways (Fig. 6c). Moreover, IL6-JAK-STAT3 signaling, TNF-α signaling via NF-κB, and inflammatory response mediated the transcriptional changes associated with Doxo treatment in MDA-MB-231 cells (Supplementary Fig. 6a). Gene set enrichment analysis (GSEA) further indicated a significant correlation between gene sets corresponding to immune-related phenotypes, such as interferon-γ response signaling, TNF-α signaling via NF-κB, and inflammatory response signaling, in Doxo-treated MCF-7 cells (Supplementary Fig. 6b). Similarly, the IL6-JAK-STAT3 signaling pathway was enriched in Abe-treated MDA-MB-231 and MCF-7 cells, and inflammatory response signaling was enriched in Abe-treated MCF-7 cells (Supplementary Fig. 6c).

Since the SASP is a characteristic phenotype of cellular senescence, we constructed the protein– protein interaction (PPI) network using STRING to determine which molecules play a crucial role in the immune and inflammatory responses under therapy-induced senescence. As shown in Fig. 6d, the commonly upregulated genes under therapy-induced senescence in the two breast cancer cell lines clustered into two distinct functional sets: senescence-related genes and immunomodulatory-related genes. These results revealed that therapy-induced cellular senescence remarkably enhanced immunomodulatory-related genes. In particular, *IL6, IL8, IL1A, IL1B, CCL2, CXCL8, NFKB1*, and *TGFB1* showed strong interaction scores and node degrees in the set of immunoregulatory-related genes. These results suggested that members of the STAT molecular superfamily, especially STAT3, likely play a significant role in SASP reprogramming.

Further, the heatmap demonstrated consistently increased immunomodulatory-related genes in therapy-induced MDA-MB-231 and MCF-7 cells (Fig. 6e and Supplementary Table 3). To validate whether the expression levels of these genes were considerably elevated under the experimental conditions, we selected the genes encoding cytokines with relatively high node degrees in the STRING network for validation analysis by RT-qPCR. Indeed, the mRNA expression levels of *CXCL2, TGFB1, CCL2*, and *IL1B* were remarkably increased in breast cancer cell lines after treatment with the therapeutic drugs (Fig. 6f and g). Together with the finding that the *IL6* expression level significantly increased in therapy-induced senescent cells, these results indicated that therapy-induced senescence in breast cancer cells triggered the expression of pro-inflammatory molecules via SASP reprogramming, which evoked a profound alteration in the transcriptional profile of genes related to inflammatory and immune responses.

## Discussion

Dysregulation in pH status has been observed in senescent human fibroblast cells due to metabolic reprogramming^45^. However, the mechanism underlying the disturbance of pH_i_ homeostasis caused by therapy-induced senescence in breast cancer is not fully understood. Here, we found that Doxo and Abe induced the expression of p21 and p16, resulting in cellular senescence in breast cancer cell lines, which is consistent with previous reports^46–50^. Moreover, we demonstrated that these therapy-induced changes in senescent cells were accompanied by significant downregulation of *ATP6AP2* expression, leading to changes in pH levels that impaired lysosome function and altered the immune response at the molecular level. We also discovered that Doxo and Abe, two commonly used breast cancer drugs, caused cell cycle arrest and reduced cell proliferation through the upregulated expression of certain genes contributing to senescence. Additionally, senescence-driven SASP reprogramming occurred, which altered inflammation and immune transcriptional profiles. Together, these findings suggest that ATP6AP2 plays a role in pH regulation during therapy-induced senescence and that altered pH homeostasis may be associated with immune changes in senescent cancer cells.

We further determined that therapy-induced senescence caused intracellular acidification and lysosomal alkalinization. Based on RNA-seq analysis of senescent cells compared to control (untreated) breast cancer cells, we found that the genes with downregulated expression in senescent cells were enriched in pH_i_ reduction- and lysosome/phagosome-related pathways. We verified the intracellular acidification in senescent cells through pHrodo Green AM staining and the lysosomal alkalinization through LysoTracker staining. These findings support the idea that therapy-induced senescence leads to changes in pH_i_ homeostasis and lysosomal function^51^.

V-ATPase is a protein complex that plays a role in regulating pH in cells, maintaining pH homeostasis within cells and intracellular organelles, which is essential for lysosomal function^52,53^. Dysregulation of V-ATPase function can lead to pH abnormalities and lysosomal dysfunction^54,55^. We demonstrated that V-ATPase activity was impaired in senescent cells (cells that have ceased dividing) induced by therapeutic drugs and that expression of the ATP6AP2 subunit was consistently downregulated in these cells. Since Baf A1 suppresses lysosomal and autophagic lysosomal functions by blocking V-ATPase activity^56,57^, we further confirmed the effect by treatment of Baf A1, a compound that disrupts V-ATPase activity, which led to more substantial changes in pH in senescent breast cancer cells than treatment with the therapeutic drugs. Together, these results reveal that ATP6AP2 suppression may play a role in the alteration of pH in senescent cells, potentially due to a lack of proton pumping, causing the accumulation of protons in the cytosol with consequent changes in pH_i_ and pH_L_. However, elucidation of the specific mechanism by which ATP6AP2 expression is downregulated in anti-cancer drug-induced senescent cells requires further investigation.

Furthermore, our study demonstrated that activation of multiple genes associated with the SASP occurs in therapy-induced senescent breast cancer cells. GSEA demonstrated that inflammatory- and immune-related pathways were significantly enriched in therapy-induced senescent cells. These findings are consistent with previous studies suggesting that senescent cells actively secrete SASP-related molecules that drive inflammatory responses and modulate the immune status^58,59^. Interestingly, we found upregulation of genes in the PD-L1 pathway in therapy-induced senescent cells, which aligns with previous studies indicating PD-L1 accumulation in senescent cells^60,61^. This may be a potential mechanism for tumor recurrence and immune evasion following treatment with therapeutic drugs. Additionally, we discovered enrichment of STAT3 expression and related pathways, supporting the relationship between senescence bypass and the immune regulation phenotype^62–64^. However, further investigation is required to fully understand the mechanisms by which senescent cells affect immune regulation.

In conclusion, our findings demonstrate that attenuated ATP6AP2 expression triggers intracellular acidification and lysosomal alkalinization in senescent breast cancer cells exposed to therapeutic drugs. The increased pH_L_ then results in lysosomal impairment. Additionally, SASP reprogramming in senescent cells upregulates multiple molecules that alter inflammatory and immune response profiles. However, the direct mechanism by which ATP6AP2 is attenuated in senescent cells remains to be determined. Further research is also required to understand the detailed mechanisms of pH dysregulation and SASP reprogramming in senescent cells and to identify the manner in which dysregulation of intracellular pH homeostasis induced by senescence leads to altered immune responses. The elucidation of these underlying mechanisms could potentially inform strategies to improve therapeutic efficacy in breast cancer.

## Methods

### Cell lines and cell culture

MDA-MB-231 (a human triple-negative breast cancer cell line) and MCF-7 (a human luminal A subtype breast cancer cell line) cells were obtained from the American Type Culture Collection (Manassas, VA, US). The cells were maintained in Dulbecco’s modified Eagle’s medium (DMEM; Nacalai Tesque, Kyoto, Japan) containing 10% fetal bovine serum (FBS; Gibco, Life Technologies, Carlsbad, CA, USA) and 0.01% penicillin/streptomycin (Wako, Osaka, Japan). Both cell lines were cultured at 37°C in a 5% v/v CO_2_ atmosphere and used for experiments at early passages (<10 passages). Both cell lines were confirmed to be negative for *Mycoplasma* contamination.

### *In vitro* senescence induction

Doxo (Sigma-Aldrich, Darmstadt, Germany) was dissolved in distilled water, whereas Abe (Verzenio, Eli Lilly, USA) was dissolved in ethanol prior to use in the experiments. MDA-MB-231 and MCF-7 cells were cultured in serum-deprived (2% FBS) conditions for 12 h. The cells were then treated with Doxo at 100 nM (the concentration determined to not induce severe cell death) and with Abe at a concentration gradient (125 nM, 250 nM, 500 nM, 1000 nM) for the initial 24 h (1st dose), respectively. The culture medium was then replaced with normal medium (without Doxo or Abe) for 48 h. Subsequently, the cells were re-exposed to the two drugs separately at the above concentrations and collected at four time points (2nd dose) for the following experiments.

### Cell proliferation assay

Cell proliferation was evaluated using the CCK-8 assay kit (Dojindo, Kumamoto, Japan) according to the manufacturer’s protocols. The cells were seeded in 96-well plates at 3000 cells/well and allowed to attach overnight. The cells were then treated with the therapeutic drugs as described above. After collecting the cells at different time points (from 24 h to 96 h), 200 μL of medium containing CCK-8 reagent was added to each well and the cells were further incubated at 37°C for 3 h. The absorbance values were measured by the SpectraMax 340PC 384 Microplate Reader (Molecular Devices, Tokyo, Japan) at a wavelength of 450 nm.

### Cell cycle assay

The cell cycle assay was performed with the PI/RNase Staining Buffer Kit (BD Biosciences, San Jose, CA, USA). A total of 3 × 10^5^ cells were seeded into each well of 6-well plates and allowed to attach overnight. The cells were harvested after treatment with the therapeutic drugs, followed by washing in phosphate-buffered saline and fixation using 70% (v/v) ice-cold ethanol for 1 h (or at 4°C overnight). The cells were then incubated with propidium iodide staining solution for 15 min at room temperature before analysis.

The cell cycle distribution was detected immediately by flow cytometry using the BD LSRFortessa Cell Analyzer (BD Biosciences, San Jose, CA, USA). Data were collected using BD FACSDiva Software (v8.0.1, BD Biosciences) and further analyzed by ModFit LT 5.0 software (Verify Software House, Topsham, ME, USA).

### SA-β-Gal assay

Cellular senescence was determined by the Senescence β-Galactosidase Staining Kit (Cell Signaling Technology, Boston, MA, USA) according to the manufacturer’s instructions. A total of 3 × 10^5^ cells were seeded into each well of a 6-well plate and treated with the therapeutic drugs as described above. The cells were then harvested, fixed with a fixative solution, and incubated with the β-Gal staining solution in a dry incubator without CO_2_ at 37°C overnight. After incubation, the cells were observed using a KEYENCE BZ-X800 fluorescence microscope (KEYENCE, Osaka, Japan). Quantification of β-Gal–positive senescent cells was performed using Image J software (v1.52, National Institutes of Health, Bethesda, MD, USA).

### RT-qPCR

Cells were treated with TRIzol reagent (Invitrogen, Carlsbad, CA, USA) and total RNA was extracted by RNeasy Mini Kit (QIAGEN Sciences, Germantown, MD, USA) following the manufacturer’s instructions. cDNA was synthesized by a reverse transcriptase reaction with 500 ng of total RNA using the Transcriptor First Strand cDNA Synthesis Kit (Roche, Basal, Switzerland), and was used as a template for qPCR with LightCycler 480 SYBR Green I Master (Roche, Basal, Switzerland) on a StepOnePlus Real-Time PCR System (Applied Biosystems, Foster City, CA, USA). Gene expression levels were normalized to the level of the endogenous control gene *ACTB* using an adjusted 2^−ΔΔCt^ method. The sequences of the primers for RT-qPCR assays are provided in Supplementary Table 4.

### Gene expression profiling

Total RNA was isolated using RNeasy Mini Kit (QIAGEN Sciences) following the manufacturer’s instructions. The RNA quality was assessed on a Nanodrop DS-11 spectrophotometer (DeNovix, Wilmington, NC, USA) to ensure a 260:280 nm ratio ≥ 2.0, and an RNA integrity number ≥ 9 was assessed by TapeStation RNA Screen Tape (Agilent). RNA-seq analysis was performed with the transcriptome for targeted next-generation sequencing by Macrogen (Tokyo, Japan). Total RNA (1 μg) was enriched with polyA + RNA, and sequencing libraries were sequenced with the TruSeq stranded mRNA Library on a NovaSeq6000 platform (Illumina, San Diego, CA, USA). RNA-seq data were checked for quality by FastQC software (v0.11.9). Sequences were aligned to the human reference genome (GRch38/hg38) with HISAT2(v2.2.1)^65^, and sequence reads were assigned to reference genomic features by FeatureCounts^66^. Computational analysis of RNA-seq data was performed by Galaxy (v22.05.1, https://usegalaxy.org/). Gene counts were scaled and normalized into transcripts per kilobase million (TPM) units. The TPM values of each gene were used to calculate the fold change (FC) and the corresponding *p*-values. Significant DEGs were identified according to the criteria of log2 FC ≥ 1.0 or log2 FC ≤ –1.0 and *p-*value ≤ 0.05.

### Functional enrichment analysis

DEGs were subjected to GO and KEGG pathway enrichment analyses using the Database for Annotation, Visualization and Integrated Discovery (DAVID) functional annotation tool^67^. Enriched GO terms and pathways were selected based on the threshold of a false discovery rate (FDR) ≤ 0.05. The PPI network was constructed with STRING (v11.5, https://string-db.org/) according to interaction score confidence (0.700), and the edges represent both functional and physical associations among proteins. The PPI network was analyzed and visualized by Cytoscape software (v3.9.1).

### GSEA

GSEA was conducted using the GSEA (v4.3.2) desktop tool^68^. Hallmark collections were acquired from the Molecular Signatures Database (MSigDB, v2022.1)^69^. Gene expression levels of therapy-treated senescent cells versus control cells were used to generate the ranked list file. A total of 1000 permutations were applied to determine the significance of the enrichment for the gene sets. The enriched phenotypes were considered significant with a *p*-value < 0.01 and FDR < 0.25, and a metric for ranking the genes was obtained by the “Signal to noise” method.

### Baf A1 treatment

Baf A1 (Bioviotica, Dransfeld, Germany) was dissolved in dimethyl sulfoxide to obtain a 20 µM stock solution. For Baf A1 treatment, the cells were cultured in the complete medium containing 200 nM Baf A1 for 3 h before pH detection.

### Measurement of pH_i_

The pH_i_ was determined using pHrodo Green AM Intracellular pH Indicator (Thermo Fisher Scientific, Waltham, MA, USA). Cells grown in DMEM (supplemented with 10% FBS) were seeded in 96-well black plates (5000 cells/well). After drug treatment, the cells were washed with Live Cell Imaging Solution (LCIS; Thermo Fisher Scientific) and labeled with pHrodo Green AM dye for 30 min at 37°C. After washing with LCIS, cell fluorescence was detected under the KEYENCE BZ-X800 fluorescence microscope (KEYENCE, Osaka, Japan) and analyzed by the SpectraMax GEMINI EM spectrofluorometer (Molecular Devices, Tokyo, Japan) at excitation (Ex) and Em wavelengths of 509 nm and 533 nm, respectively. A pH_i_ standard curve was constructed by the Intracellular pH Calibration Buffer Kit (Thermo Fisher Scientific). Cells were treated with different pH calibration solutions (pH 4.5, 5.5, 6.5, and 7.5), and then the fluorescence intensity was measured to fit a linear trendline to obtain the pH standard curve.

### pH_L_ detection

The pH_L_ was detected using the ratiometric lysosomal pH dye LysoSensor Yellow/Blue DND-160 (Thermo Fisher Scientific). Cells were seeded in 96-well black plates at 5000 cells/well. After drug treatment, the cells were washed with LCIS and incubated with 2 µM LysoSensor Yellow/Blue for 5 min at 37°C under 5% CO_2_. After washing with LCIS, cell fluorescence was detected under the KEYENCE BZ-X800 fluorescence microscope and measured by the Multilabel Plate Reader ARVO X5 (PerkinElmer, Waltham, MA, USA) at Em 440 nm and 535 nm in response to Ex at 340 and 380 nm, respectively. The ratio of light emitted with 340 and 380 nm excitation was used to determine the pH_L_ status.

### Statistics

All results are presented as the mean ± standard deviation (SD) of three independent experiments. Prior to analysis, a normality test was performed on all collected data. A two-tailed unpaired Student’s t test was used for pairwise comparisons, whereas multiple comparisons were evaluated using one-way or two-way analysis of variance, followed by Dunnett’s, Sidak’s, or Tukey’s multiple comparisons test. Correlation analysis of two independent samples was performed using Pearson’s correlation coefficient. Two-tailed *p-* values ≤ 0.05 were considered statistically significant. Data analyses were conducted using GraphPad Prism software (v9.4.1, GraphPad, La Jolla, CA, USA).

### Data Availability

The RNA-seq data generated in this study have been deposited in the Gene Expression Omnibus (GEO) (URL: https://www.ncbi.nlm.nih.gov/geo/) database under accession code GSE222984. The remaining data are available within the article and Supplementary Data files 1-4. In addition, any other data that support the findings of this study can be made available on request from the corresponding author.

## Notes

### Competing Interest Statement

K.K.: grants from TERUMO, Astellas, Eli Lilly, Kyoto Breast Cancer Research Network; consulting fees from Becton Dickinson Japan; honoraria from Eisai, Chugai, and Takeda. M.T.: grants from Chugai, Takeda, Pfizer, Kyowa-Kirin, Taiho, JBCRG Associates, Eisai, Eli Lilly, Daiichi-Sankyo, AstraZeneca, Astellas, Shimadzu, Yakult, Nippon Kayaku, AFI Technology, Luxonus, Shionogi, and GL Science; honoraria from Chugai, Takeda, Pfizer, Kyowa-Kirin, Taiho, Eisai, Daiichi-Sankyo, AstraZeneca, Eli Lilly, MSD, Exact Science, Novartis, Konica Minolta, Shimadzu, Yakult, and Nippon Kayaku; advisory board of Kyowa-Kirin, Daiichi-Sankyo, Eli Lilly, Konica Minolta, BMS, Athenex Oncology, Bertis, Terumo, Kansai Medical Net; board of directors of JBCRG Associates, KBCRN, OOTR; Associate Editor of the British Journal of Cancer, Scientific Reports, Breast Cancer Research and Treatment, Cancer Science, Frontiers in Women's Cancer, Asian Journal of Surgery, Asian Journal of Breast Surgery; deputy editor of International Journal of Oncology.

