## Supplementary Information for "Senescence triggers intracellular acidification and lysosomal alkalinization via ATP6AP2 attenuation in breast cancer cells"

Supplementary Fig. 1 is related to Fig. 1

Supplementary Fig. 2 is related to Fig. 2

Supplementary Fig. 3 is related to Fig. 3

Supplementary Fig. 4 is related to Fig. 4

Supplementary Fig. 5 is related to Fig. 5

Supplementary Fig. 6 is related to Fig. 6

Supplementary Data 1 is related to Fig. 3 and Fig. 6

Supplementary Data 2 is related to Fig. 3 and Fig. 6

Supplementary Data 3 is related to Fig. 3 and Fig. 6

Supplementary Data 4 is related to Fig. 3 and Fig. 6

Supplementary Data 5 is related to Fig. 1 and Supplementary Fig. 1

Supplementary Table 1 is related to Fig. 3

Supplementary Table 2 is related to Supplementary Fig. 5

Supplementary Table 3. is related to Fig. 6

Supplementary Table 4. For all Figs

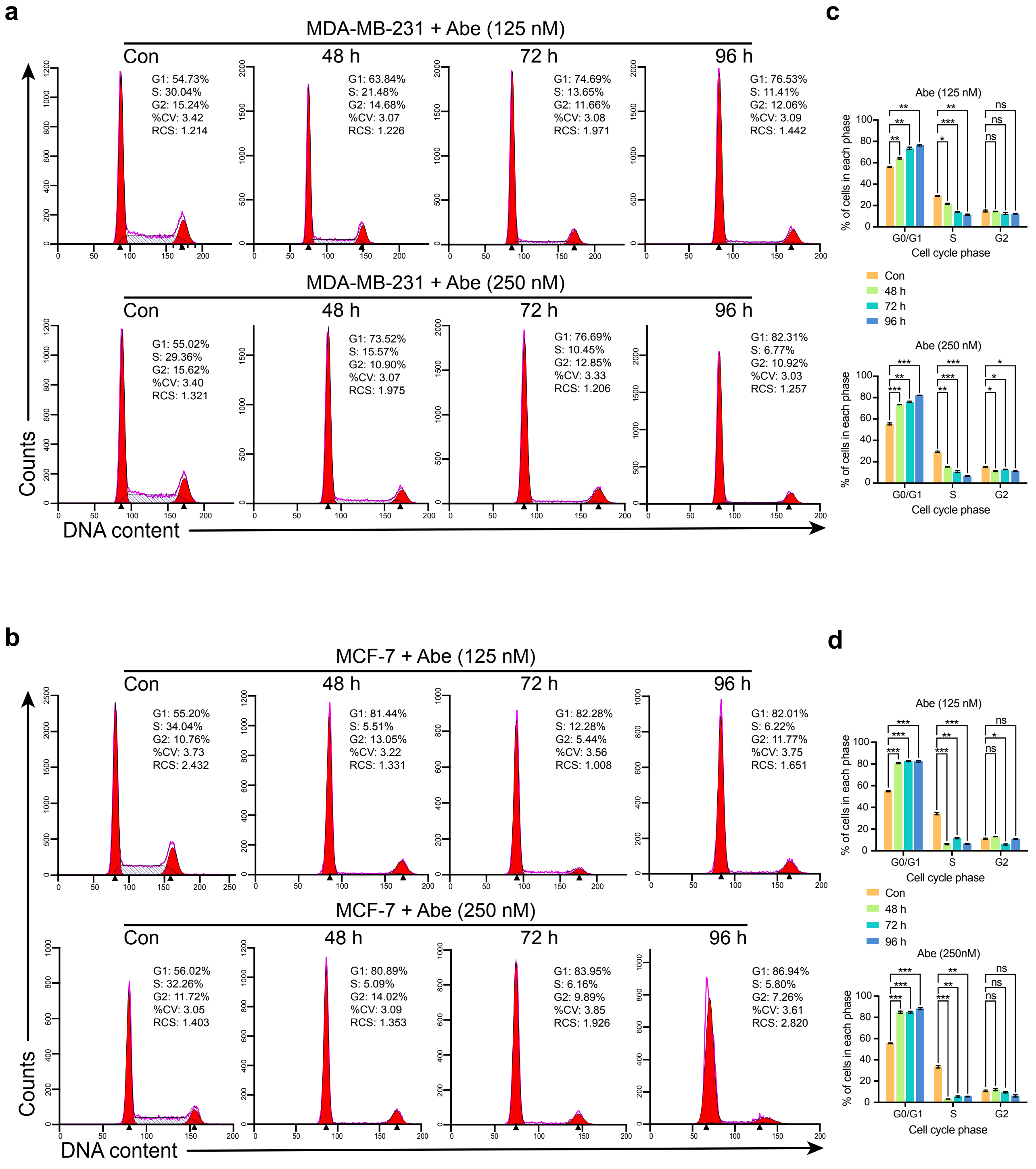


**Supplementary Fig. 1. Abemaciclib induces G1 cell cycle arrest in breast cancer cells.**

(**a** and **b**) Representative images of the cell cycle analysis using propidium iodine (PI) detection by flow cytometry in Abe (125 nM or 250 nM)-treated MDA-MB-231 (**a**) and MCF-7 (**b**).

(**c** and **d**) Quantification of the percentage in each cell cycle phase, respectively.

Data are shown as means ± SD of three independent experiments. Two-way ANOVA with Dunnett's multiple-comparisons test (**c** and **d**) was performed. ns, not significant. *, $P < 0.05$; **, $P < 0.01$; ***, $P < 0.001$

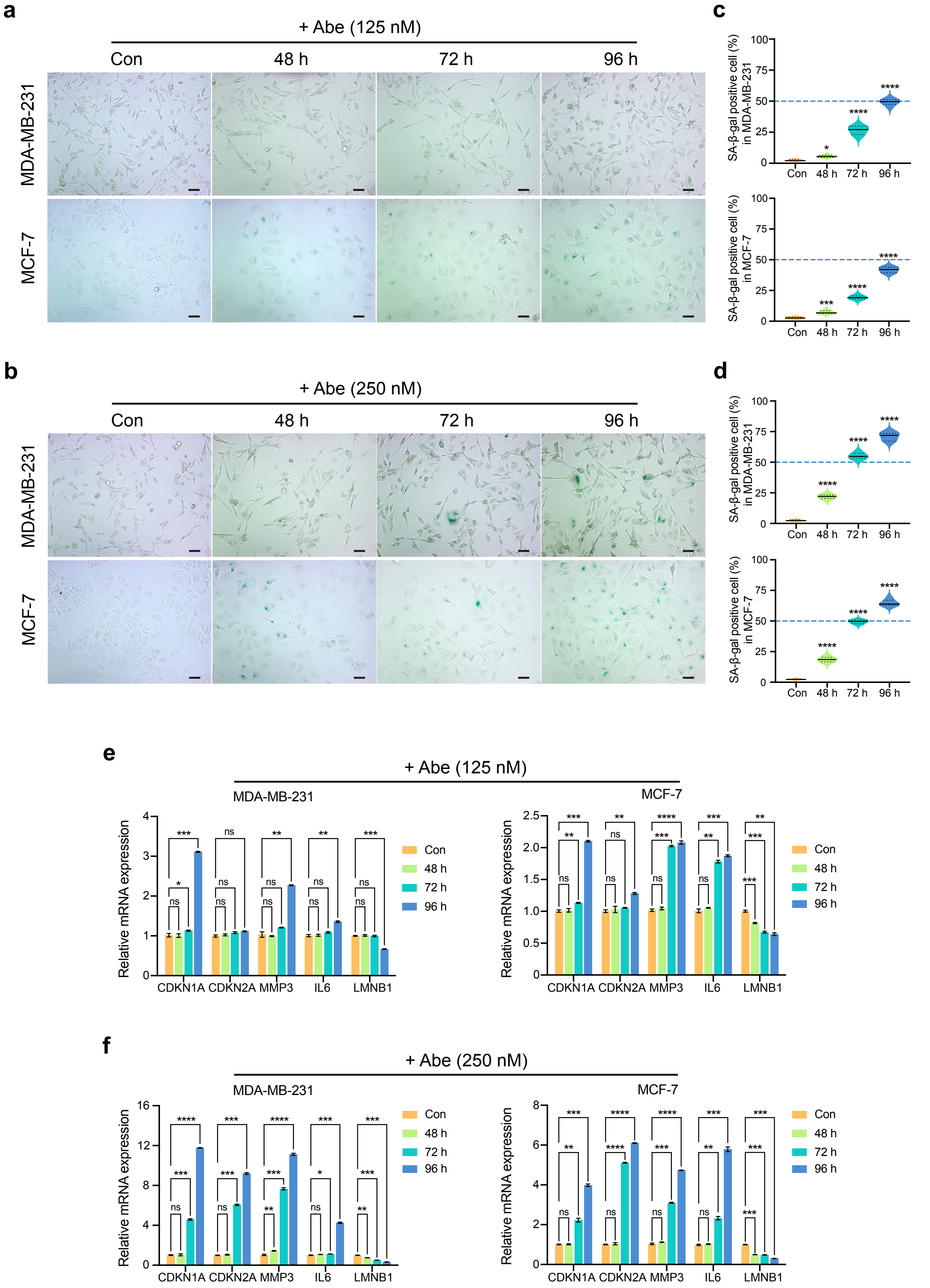
a
+ Abe (125 nM)
Con
48 h
72 h
96 h
MDA-MB-231
MCF-7
b
+ Abe (250 nM)
Con
48 h
72 h
96 h
MDA-MB-231
MCF-7
c
SA-β-gal positive cell (%) in MDA-MB-231
SA-β-gal positive cell (%) in MCF-7
d
SA-β-gal positive cell (%) in MDA-MB-231
SA-β-gal positive cell (%) in MCF-7
Con
48 h
72 h
96 h
e
+ Abe (125 nM)
MDA-MB-231
MCF-7
Relative mRNA expression
CDKN1A
CDKN2A
MMP3
IL6
LMNB1
Con
48 h
72 h
96 h
f
+ Abe (250 nM)
MDA-MB-231
MCF-7
Relative mRNA expression
CDKN1A
CDKN2A
MMP3
IL6
LMNB1
Con
48 h
72 h
96 h

**Supplementary Fig. 2. Senescent phenotypes are expressed by Abemaciclib-treated breast cancer cells.**

**(a** and **b)** Representative SA-β-gal staining in Abe (125 nM)-treated (**a**) and Abe (250 nM)-treated (**b**) breast cancer cells.

**(c** and **d)** Quantitative analysis of the percentage of SA-β-gal positive cells, respectively. At least six separate fields of view were taken.

**(e** and **f)** RT-qPCR analyzed mRNA relative expression levels of senescence-related genes (*CDKN1A, CDKN21, MMP3, IL6, LMNB1*), respectively.

Scale bars represent 50 μm. Data are shown as means ± SD of three independent experiments. One-way ANOVA with Dunnett's multiple comparisons test (**c** and **d**) and Two-way ANOVA with Dunnett's multiple-comparisons test (**e** and **f**) were performed. ns, not significant. *, $P < 0.05$; **, $P < 0.01$; ***, $P < 0.001$; ****, $P < 0.0001$.

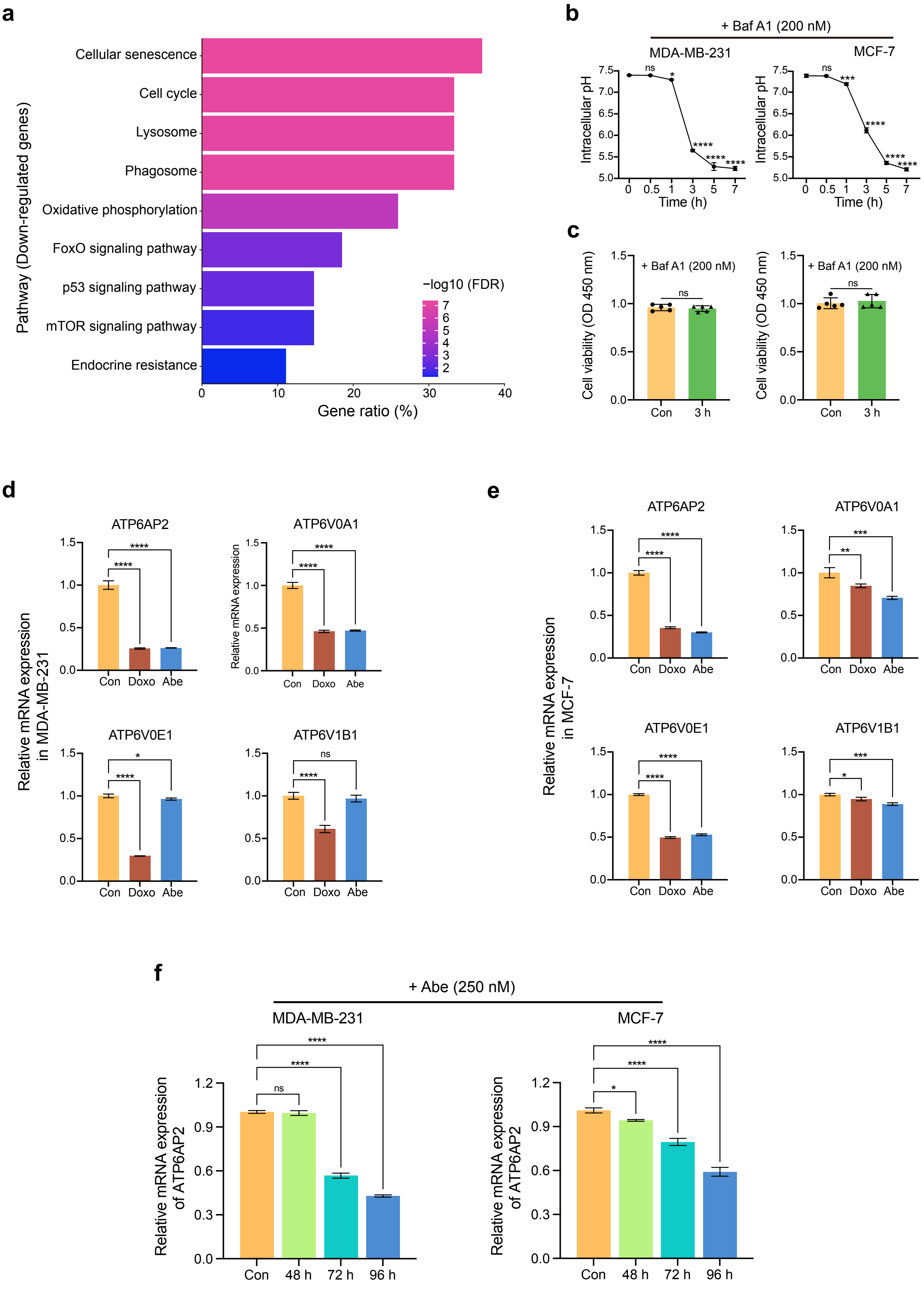
a
Pathway (Down-regulated genes)
Cellular senescence
Cell cycle
Lysosome
Phagosome
Oxidative phosphorylation
FoxO signaling pathway
p53 signaling pathway
mTOR signaling pathway
Endocrine resistance
−log10 (FDR)
Gene ratio (%)
b
+ Baf A1 (200 nM)
MDA-MB-231
MCF-7
Intracellular pH
Time (h)
c
Cell viability (OD 450 nm)
+ Baf A1 (200 nM)
Con
3 h
d
Relative mRNA expression in MDA-MB-231
ATP6AP2
ATP6V0A1
ATP6V0E1
ATP6V1B1
Con Doxo Abe
e
Relative mRNA expression in MCF-7
ATP6AP2
ATP6V0A1
ATP6V0E1
ATP6V1B1
Con Doxo Abe
f
+ Abe (250 nM)
MDA-MB-231
MCF-7
Relative mRNA expression of ATP6AP2
Con 48 h 72 h 96 h

**Supplementary Fig. 3. ATP6AP2 downregulates in senescent cells facilitated by therapeutic drugs.**

**(a)** Pathway enrichment analysis of co-downregulated genes (n = 29). All represented pathway terms with FDR < 0.05.

**(b)** Baf A1 (200 nM) regulates $pH_i$ in a time-dependent manner.

**(c)** The CCK-8 assay determined cell viability in breast cancer cells treated with Baf A1 (200 nM) for three hours.

**(d** and **e)** RT-qPCR validated the relative expression levels of V-ATPase subtypes, respectively.

**(f)** RT-qPCR analyzed mRNA relative expression levels of *ATP6AP2* in breast cancer cells treated with 250 nM Abe.

Data are shown as means ± SD of three independent experiments. One-way ANOVA with Dunnett's multiple-comparisons test (**b**, **c**, and **d**) and Unpaired two-tailed Student's t test (**e**) were performed. ns, not significant. *, *P* < 0.05; **, *P* < 0.01; ***, *P* < 0.001; ****, *P* < 0.0001.

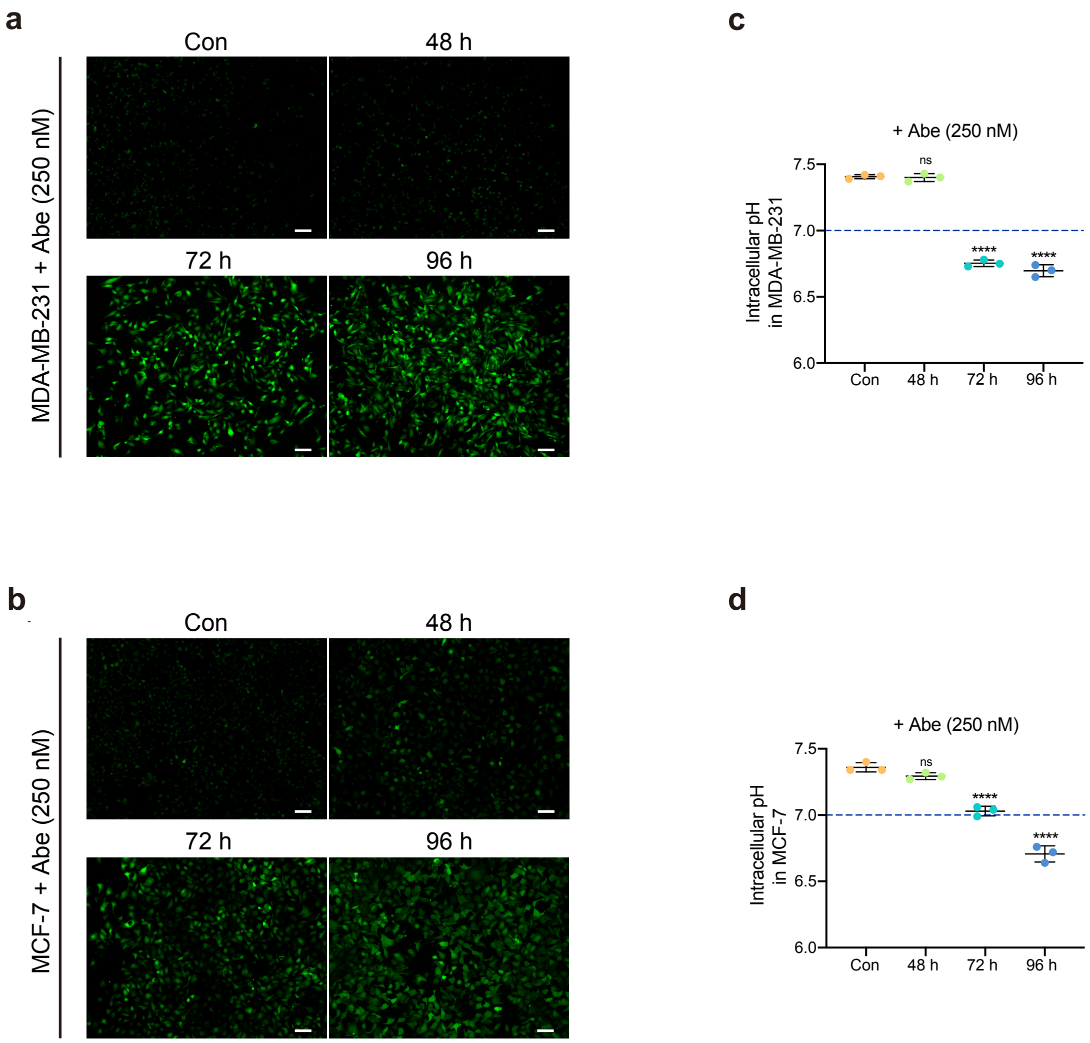


**Supplementary Fig. 4. Abemaciclib induces intracellular pH decrease in breast cancer cells.**

(**a** and **b**) Representative fluorescent images of Abe (250 nM)-treated MDA-MB-231 (**a**) and MCF-7 (**b**) stained with pHrodo Green AM.

(**c** and **d**) Quantitative analysis of the $pH_i$ in MDA-MB-231 (**c**) and MCF-7 cells (**d**).

Scale bars represent 50 μm. Data are shown as means ± SD of three independent experiments. One-way ANOVA with Dunnett's multiple-comparisons test (**c** and **d**) were performed. ns, not significant. ****, $P < 0.0001$.

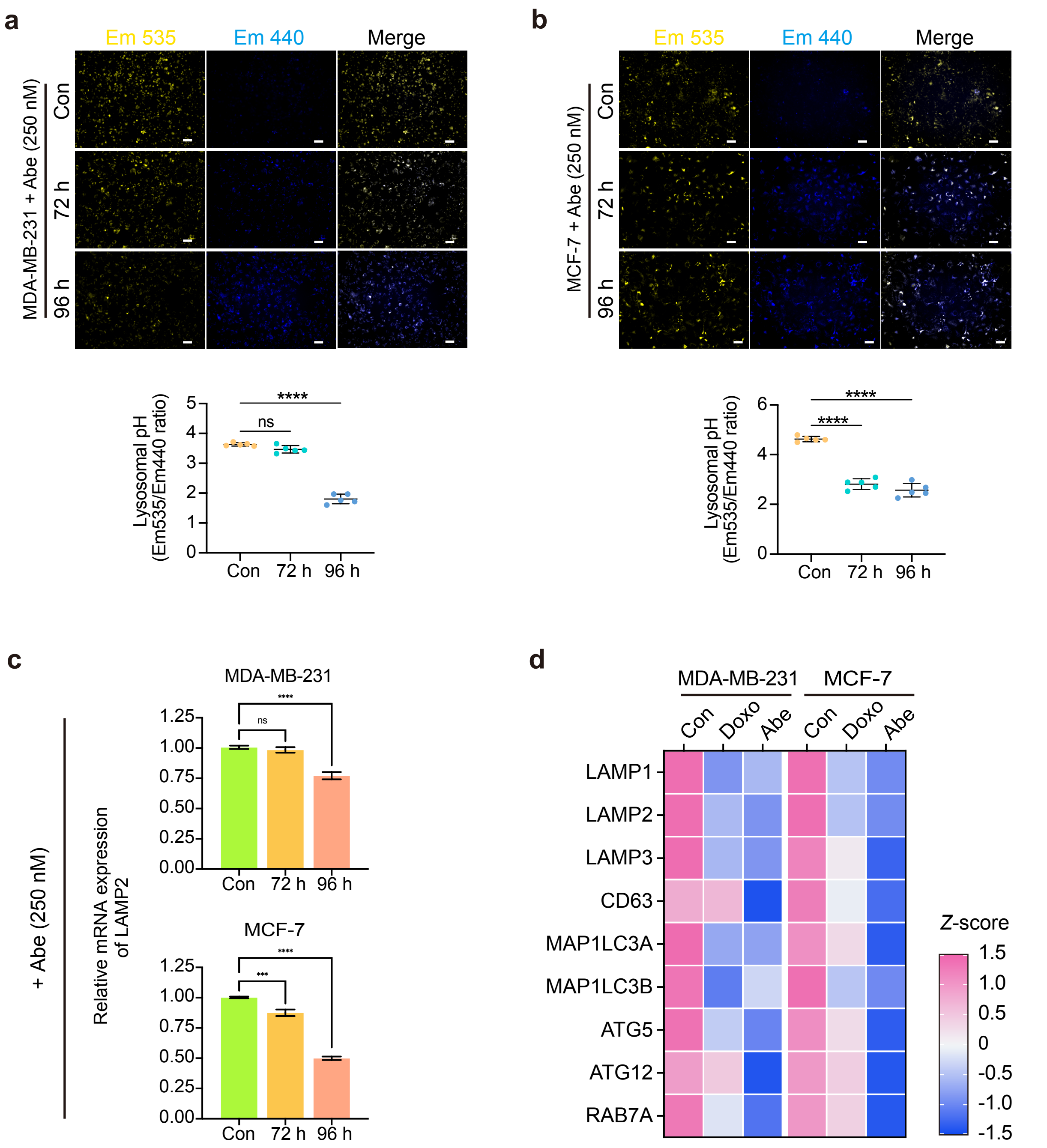


**Supplementary Fig. 5. Therapeutic drugs increase lysosomal pH and trigger lysosomal dysfunction.**

(**a** and **b**) Representative fluorescent images of Abe (250 nM)-treated MDA-MB-231 (**a** upper) and MCF-7 (**b** upper) stained with LysoSensor Yellow/Blue-DND-160 to assess $pH_L$ status. Em 535: yellow, Em440: blue. Quantitative analysis of the $pH_L$ status (lower panel).

**(c)** Heatmap represents the standardized mRNA abundance values (*z*-score) of the lysosomal function-related genes.

**(d)** RT-qPCR analyzed the mRNA relative expression levels of *LAMP2*.

Scale bars represent 50 μm. Data are shown as means ± SD of three independent experiments. One-way ANOVA with Dunnett's multiple-comparisons test (**a**, **b** lower and **d**) was performed. ns, not significant. ***, $P < 0.001$; ****, $P < 0.0001$.

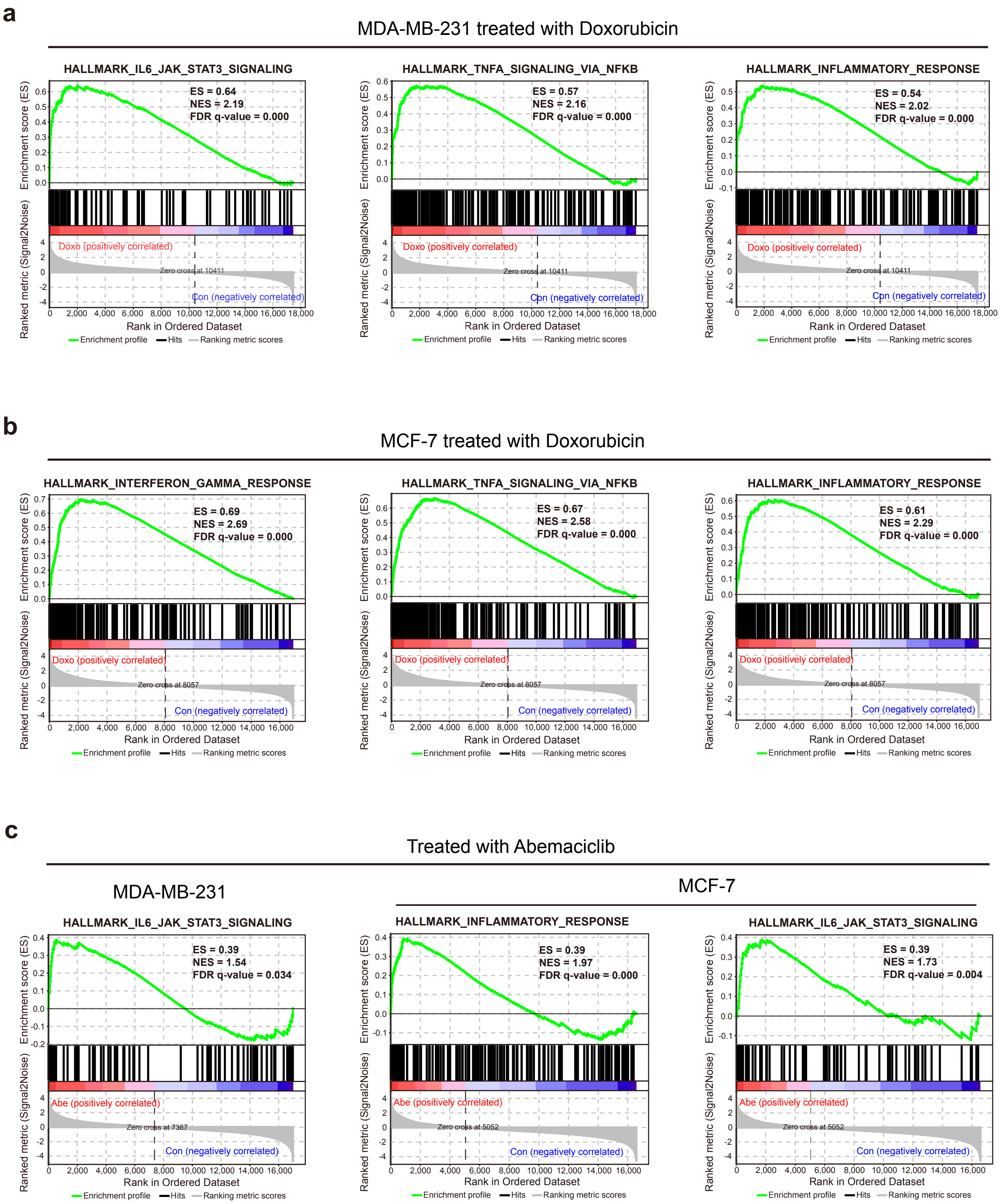
a
MDA-MB-231 treated with Doxorubicin
HALLMARK_IL6_JAK_STAT3_SIGNALING
ES = 0.64
NES = 2.19
FDR q-value = 0.000
HALLMARK_TNFA_SIGNALING_VIA_NFKB
ES = 0.57
NES = 2.16
FDR q-value = 0.000
HALLMARK_INFLAMMATORY_RESPONSE
ES = 0.54
NES = 2.02
FDR q-value = 0.000
Doxo (positively correlated)
Zero cross at 10411
Con (negatively correlated)
b
MCF-7 treated with Doxorubicin
HALLMARK_INTERFERON_GAMMA_RESPONSE
ES = 0.69
NES = 2.69
FDR q-value = 0.000
HALLMARK_TNFA_SIGNALING_VIA_NFKB
ES = 0.67
NES = 2.58
FDR q-value = 0.000
HALLMARK_INFLAMMATORY_RESPONSE
ES = 0.61
NES = 2.29
FDR q-value = 0.000
Doxo (positively correlated)
Zero cross at 8057
Con (negatively correlated)
c
Treated with Abemaciclib
MDA-MB-231
MCF-7
HALLMARK_IL6_JAK_STAT3_SIGNALING
ES = 0.39
NES = 1.54
FDR q-value = 0.034
Zero cross at 7367
HALLMARK_INFLAMMATORY_RESPONSE
ES = 0.39
NES = 1.97
FDR q-value = 0.000
Zero cross at 5052
HALLMARK_IL6_JAK_STAT3_SIGNALING
ES = 0.39
NES = 1.73
FDR q-value = 0.004
Zero cross at 5052
Abe (positively correlated)
Con (negatively correlated)
Enrichment score (ES)
Ranked metric (Signal2Noise)
Rank in Ordered Dataset
Enrichment profile
Hits
Ranking metric scores

**Supplementary Fig. 6. Inflammatory and immune response pathways enriched in therapy-induced senescent cells.**

**(a)** GSEA enrichment analysis was performed in Doxo (100 nM)-treated MDA-MB-231 cells.

(**b**) GSEA enrichment analysis was performed in Doxo (100 nM)-treated MCF-7 cells.

(**c**) GSEA enrichment analysis was performed in Abe (500 nM)-treated MDA-MB-231 (left panel) and MCF-7 (right panel).

Genes (vertical black lines) represented in gene sets are on the x-axis, and the y-axis represents the enrichment score (ES). The green line connects genes and ES points. The colored band shows the degree of correlation of genes with the enriched phenotype (red, positive correlation, and blue, negative correlation). FDR < 0.05 as the significance threshold.

**Supplementary Table 1**

| Group | MDA-MB-231 (TPM) | | | MCF-7 (TPM) | | |
|---|---|---|---|---|---|---|
| | 231-Con | 231-Doxo | 231-Abe | 7-Con | 7-Doxo | 7-Abe |
| **ATP6AP2** | 167.141 | 62.8521 | 65.97617 | 135.0201 | 54.24743 | 54.52637 |
| **ATP6V0A1** | 24.6106 | 11.18999 | 11.11805 | 21.62104 | 8.411995 | 6.514685 |
| **ATP6V0E1** | 491.2237 | 230.3726 | 234.0847 | 483.068 | 223.2373 | 157.2103 |
| **ATP6V1B1** | 0.837144 | 0.27683 | 0.354183 | 10.20945 | 3.942186 | 4.789519 |
| **ATP6V0E2** | 40.33683 | 19.86086 | 15.44662 | 28.86607 | 6.111483 | 8.635479 |
| **ATP6V1H** | 45.47745 | 17.56449 | 20.28437 | 39.5637 | 13.18008 | 14.08679 |
| **ATP6V1F** | 233.3758 | 109.5692 | 99.95342 | 221.4649 | 76.48275 | 85.15201 |
| **ATP6V1G1** | 139.5136 | 53.32301 | 64.6393 | 225.3341 | 110.5389 | 103.3351 |

| Group | MDA-MB-231 | | | | |
|---|---|---|---|---|---|
| | Average | SD | z-score | | |
| **ATP6AP2** | 98.65643 | 48.44271 | 1.413723 | -0.739107 | -0.674617 |
| **ATP6V0A1** | 15.63955 | 6.343562 | 1.414198 | -0.701429 | -0.71277 |
| **ATP6V0E1** | 318.5603 | 122.1009 | 1.414105 | -0.722253 | -0.691851 |
| **ATP6V1B1** | 0.489386 | 0.247922 | 1.402694 | -0.85735 | -0.545344 |
| **ATP6V0E2** | 25.21477 | 10.84371 | 1.394547 | -0.493734 | -0.900813 |
| **ATP6V1H** | 27.77544 | 12.56637 | 1.408682 | -0.812562 | -0.59612 |
| **ATP6V1F** | 147.6328 | 60.75638 | 1.411258 | -0.626495 | -0.784763 |
| **ATP6V1G1** | 85.82532 | 38.24344 | 1.403857 | -0.849879 | -0.553978 |

| Group | MCF-7 | | | | |
|---|---|---|---|---|---|
| | Average | SD | z-score | | |
| **ATP6AP2** | 81.26465 | 38.01105 | 1.414207 | -0.710773 | -0.703434 |
| **ATP6V0A1** | 12.18257 | 6.7188 | 1.404784 | -0.561198 | -0.843586 |
| **ATP6V0E1** | 287.8385 | 140.6551 | 1.388001 | -0.459288 | -0.928713 |
| **ATP6V1B1** | 6.313719 | 2.776333 | 1.403193 | -0.854196 | -0.548997 |
| **ATP6V0E2** | 14.53768 | 10.18397 | 1.406956 | -0.827398 | -0.579558 |
| **ATP6V1H** | 22.27686 | 12.22925 | 1.413566 | -0.743854 | -0.669711 |
| **ATP6V1F** | 127.6999 | 66.39627 | 1.412203 | -0.771386 | -0.640817 |
| **ATP6V1G1** | 146.4027 | 55.89035 | 1.412254 | -0.641681 | -0.770573 |

**Supplementary Table 1.** RNA-seq normalization results for representative subunits of V-ATPase in 29 co-downregulated genes.

**Supplementary Table 2**

| Group | MDA-MB-231 (TPM) | | | MCF-7 (TPM) | | |
|---|---|---|---|---|---|---|
| | 231-Con | 231-Doxo | 231-Abe | 7-Con | 7-Doxo | 7-Abe |
| **LAMP1** | 254.8058 | 75.95875 | 100.4852 | 132.6199 | 61.99424 | 42.95981 |
| **LAMP2** | 164.0145 | 79.34554 | 64.82179 | 51.30565 | 24.03746 | 17.01098 |
| **LAMP3** | 0.512301 | 0.157638 | 0.104327 | 3.160298 | 2.218661 | 0.979365 |
| **CD63** | 433.934 | 424.0611 | 256.0624 | 459.6124 | 371.0348 | 294.0614 |
| **MAP1LC3A** | 13.57517 | 6.84313 | 6.679199 | 0.590627 | 0.438305 | 0.118992 |
| **MAP1LC3B** | 6.044111 | 3.629767 | 4.43065 | 75.12739 | 62.5088 | 58.89508 |
| **ATG5** | 28.19634 | 23.09392 | 21.26248 | 28.06269 | 24.01569 | 16.14677 |
| **ATG12** | 30.48079 | 30.41372 | 30.14027 | 29.47602 | 28.84029 | 26.70154 |
| **RAB7A** | 257.5688 | 209.4985 | 176.5374 | 256.4669 | 249.5 | 229.1101 |

| Group | MDA-MB-231 | | | | |
|---|---|---|---|---|---|
| | Average | SD | z-score | | |
| **LAMP1** | 143.7499 | 79.16414 | 1.402856 | -0.856337 | -0.546519 |
| **LAMP2** | 102.7273 | 43.74033 | 1.40116 | -0.534557 | -0.866602 |
| **LAMP3** | 0.258089 | 0.181068 | 1.403961 | -0.55477 | -0.849191 |
| **CD63** | 371.3525 | 81.62199 | 0.766724 | 0.645765 | -1.412488 |
| **MAP1LC3A** | 9.032501 | 3.212853 | 1.413907 | -0.681442 | -0.732465 |
| **MAP1LC3B** | 4.701509 | 1.004088 | 1.337136 | -1.067379 | -0.269757 |
| **ATG5** | 24.18425 | 2.933852 | 1.367518 | -0.371638 | -0.995881 |
| **ATG12** | 30.34493 | 0.147285 | 0.922474 | 0.467085 | -1.389559 |
| **RAB7A** | 214.5349 | 33.27204 | 1.293394 | -0.15137 | -1.142024 |

| Group | MDA-MB-231 | | | | |
|---|---|---|---|---|---|
| | Average | SD | z-score | | |
| **LAMP1** | 79.19133 | 38.57062 | 1.385215 | -0.44586 | -0.939355 |
| **LAMP2** | 30.7847 | 14.79133 | 1.387364 | -0.456162 | -0.931202 |
| **LAMP3** | 2.119441 | 0.893122 | 1.165413 | 0.111094 | -1.276507 |
| **CD63** | 374.9029 | 67.64126 | 1.252336 | -0.057185 | -1.195151 |
| **MAP1LC3A** | 0.382642 | 0.196526 | 1.058312 | 0.283236 | -1.341549 |
| **MAP1LC3B** | 65.51043 | 6.958414 | 1.382063 | -0.431366 | -0.950697 |
| **ATG5** | 22.74172 | 4.94736 | 1.075518 | 0.257505 | -1.333023 |
| **ATG12** | 28.33928 | 1.186785 | 0.957828 | 0.422155 | -1.379982 |
| **RAB7A** | 245.0257 | 11.60785 | 0.985646 | 0.385458 | -1.371104 |

**Supplementary Table 2.** RNA-seq normalization results for lysosome-related genes in 29 co-downregulated genes.

**Supplementary Table 3**

| Group | MDA-MB-231 (TPM) | | | MCF-7 (TPM) | | |
|---|---|---|---|---|---|---|
| | 231-Con | 231-Doxo | 231-Abe | 7-Con | 7-Doxo | 7-Abe |
| **IL1A** | 1.041938 | 13.23502 | 13.70842 | 0.047229 | 0.109261 | 0.104689 |
| **IL1B** | 1.75634 | 10.90239 | 15.37293 | 0.044085 | 0.170377 | 0.116485 |
| **IL6** | 20.3943 | 168.9246 | 69.61748 | 0.039469 | 0.768743 | 0.250759 |
| **CCL2** | 1.288564 | 3.270175 | 4.423768 | 0.103827 | 0.316935 | 0.437487 |
| **CCL5** | 1.326898 | 4.118867 | 4.263787 | 1.189923 | 7.289544 | 10.12159 |
| **CXCL2** | 6.428843 | 81.91512 | 121.4013 | 0.081051 | 0.392165 | 0.342842 |
| **CXCL8** | 60.60142 | 147.2887 | 187.0628 | 0.057282 | 0.587602 | 0.346597 |
| **CXCL10** | 0.439703 | 7.147013 | 5.834653 | 0.038827 | 1.606925 | 0.135234 |
| **TGFB1** | 87.40451 | 206.7277 | 204.0008 | 10.1757 | 24.62314 | 28.43897 |
| **TGFB2** | 25.75662 | 53.76866 | 59.4323 | 5.034845 | 10.14993 | 16.38193 |
| **TNFSF10** | 55.95949 | 125.6745 | 137.2826 | 7.054577 | 25.9531 | 42.81299 |
| **TNFSF15** | 6.08423 | 41.41327 | 39.10266 | 0.096319 | 0.803825 | 0.1533 |
| **MMP3** | 0.086773 | 3.095388 | 2.991923 | 0.03592 | 0.3155 | 0.161222 |
| **CGAS** | 26.09697 | 60.89907 | 123.4569 | 8.696542 | 25.9113 | 24.04705 |
| **STING1** | 36.83351 | 87.00439 | 95.9179 | 1.630873 | 4.635319 | 5.433757 |
| **STAT3** | 49.17544 | 118.2252 | 123.8268 | 47.42352 | 111.4084 | 111.984 |
| **NFKB1** | 19.29642 | 49.29644 | 41.49849 | 31.1816 | 71.24776 | 74.47357 |
| **NFKB2** | 43.92448 | 108.6238 | 96.17982 | 10.42714 | 23.78766 | 29.42708 |

| Group | MDA-MB-231 | | | | |
|---|---|---|---|---|---|
| | Average | SD | z-score | | |
| **IL1A** | 9.328459 | 5.862641 | -1.413445 | 0.666349 | 0.747096 |
| **IL1B** | 9.343889 | 5.667134 | -1.338869 | 0.275007 | 1.063861 |
| **IL6** | 86.31214 | 61.77567 | -1.067052 | 1.337298 | -0.270247 |
| **CCL2** | 2.994169 | 1.294736 | -1.317338 | 0.213175 | 1.104163 |
| **CCL5** | 3.236517 | 1.3516 | -1.412858 | 0.652818 | 0.76004 |
| **CXCL2** | 69.91507 | 47.69811 | -1.331001 | 0.251583 | 1.079418 |
| **CXCL8** | 131.651 | 52.79852 | -1.345674 | 0.296178 | 1.049496 |
| **CXCL10** | 4.47379 | 2.902409 | -1.38991 | 0.921036 | 0.468874 |
| **TGFB1** | 166.0443 | 55.6179 | -1.41393 | 0.73148 | 0.68245 |
| **TGFB2** | 46.31919 | 14.72263 | -1.396664 | 0.505987 | 0.890677 |
| **TNFSF10** | 106.3055 | 35.91405 | -1.401848 | 0.539314 | 0.862533 |
| **TNFSF15** | 28.86672 | 16.13725 | -1.411795 | 0.77749 | 0.634305 |
| **MMP3** | 2.058028 | 1.394528 | -1.413565 | 0.743879 | 0.669686 |
| **CGAS** | 70.151 | 40.28184 | -1.093645 | -0.22968 | 1.323325 |
| **STING1** | 73.25193 | 26.00755 | -1.400302 | 0.528787 | 0.871515 |
| **STAT3** | 97.07582 | 33.94779 | -1.411001 | 0.622998 | 0.788004 |

| | | | | | |
|---|---|---|---|---|---|
| **NFKB1** | 36.69712 | 12.70932 | -1.369129 | 0.991345 | 0.377783 |
| **NFKB2** | 82.90938 | 28.0307 | -1.390793 | 0.917368 | 0.473425 |

| **Group** | **MCF-7** | | | | |
|---|---|---|---|---|---|
| | Average | SD | z-score | | |
| **IL1A** | 0.08706 | 0.028226 | -1.411119 | 0.786543 | 0.624576 |
| **IL1B** | 0.110316 | 0.051742 | -1.279997 | 1.160772 | 0.119225 |
| **IL6** | 0.35299 | 0.306375 | -1.023324 | 1.357006 | -0.333682 |
| **CCL2** | 0.286083 | 0.137952 | -1.321155 | 0.223643 | 1.097512 |
| **CCL5** | 6.200352 | 3.726788 | -1.344436 | 0.29226 | 1.052176 |
| **CXCL2** | 0.272019 | 0.136528 | -1.398748 | 0.880007 | 0.518741 |
| **CXCL8** | 0.330494 | 0.216801 | -1.260192 | 1.185917 | 0.074276 |
| **CXCL10** | 0.593662 | 0.717565 | -0.773219 | 1.412085 | -0.638865 |
| **TGFB1** | 21.07927 | 7.865789 | -1.386201 | 0.450542 | 0.935659 |
| **TGFB2** | 10.52224 | 4.639904 | -1.182652 | -0.08024 | 1.262892 |
| **TNFSF10** | 25.27356 | 14.60622 | -1.247344 | 0.046524 | 1.20082 |
| **TNFSF15** | 0.351148 | 0.320935 | -0.79402 | 1.410494 | -0.616473 |
| **MMP3** | 0.170881 | 0.114342 | -1.180322 | 1.264794 | -0.084473 |
| **CGAS** | 19.55163 | 7.713348 | -1.407312 | 0.824502 | 0.58281 |
| **STING1** | 3.899983 | 1.637278 | -1.385904 | 0.449121 | 0.936783 |
| **STAT3** | 90.27195 | 30.29933 | -1.414171 | 0.697587 | 0.716584 |
| **NFKB1** | 58.96764 | 19.69178 | -1.411047 | 0.623616 | 0.787431 |
| **NFKB2** | 21.21396 | 7.967322 | -1.353882 | 0.323031 | 1.030851 |

**Supplementary Table 1.** RNA-seq normalization results for immunomodulatory-related genes of V-ATPase in 66 co-upregulated genes.

**Supplementary Table 4**

| Gene | Forward | Reverse |
|---|---|---|
| CDKN1A | GGATGTCCGTCAGAACCC | GCTCCCAGGCGAAGTCA |
| CDKN2A | ATGGAGCCTTCGGCTGACT | GTAACTATTCGGTGCGTTGGG |
| IL6 | TAGTCCTTCCTACCCCAATTTCC | TTGGTCCTTAGCCACTCCTTC |
| IL1B | CCACAGACCTTCCAGGAGAATG | GTGCAGTTCAGTGATCGTACAGG |
| MMP3 | CTGGACTCCGACACTCTGGA | CAGGAAAGGTTCTGAAGTGACC |
| LMNB1 | GAAAAAGACAACTCTCGTCGCA | GTAAGCACTGATTTCCATGTCCA |
| ATP6AP2 | AGGCAGTGTCATTTCGTACC | GCCTTCCCTACCATATACACTC |
| ATP6V0A1 | AGGCTGAAATCGAGAACCCC | GCTCGGAACCCTTCACAGAT |
| ATP6V0E1 | GTCCTAACCGGGGAGTTATCA | AAAGAGAGGGTTGAGTTGGGC |
| ATP6V1B1 | GGCGGTCACCCGAAACTAC | GGACGATCTCCGCATACTGG |
| LAMP2 | GAAAATGCCACTTGCCTTTATGC | AGGAAAAGCCAGGTCCGAAC |
| CXCL2 | GGGCAGAAAGCTTGTCTCAA | GCTTCCTCCTTCCTTCTGGT |
| CCL2 | AGAATCACCAGCAGCAAGTGTCC | TCCTGAACCCACTTCTGCTTGG |
| TGFB1 | TACCTGAACCCGTGTTGCTCTC | GTTGCTGAGGTATCGCCAGGAA |
| ACTB | ATTGGCAATGAGCGGTTC | GGATGCCACAGGACTCCAT |

**Supplementary Table 4.** Sequences of qRT-PCR primers. Sequences are represented as 5’ to 3’.
